# Designing protein/non-protein binding interactions using a full-atom diffusion model

**DOI:** 10.64898/2026.02.02.693502

**Authors:** Kale Kundert, George M. Church

## Abstract

An unresolved challenge in the field of computational protein design is to create proteins that bind non-protein partners, e.g. DNA, RNA, and small molecules. Most machine learning (ML) algorithms for protein design can only work with systems composed entirely of amino acids, and therefore cannot be directly applied to this task. The few algorithms that accommodate non-proteins still represent amino acids differently than other molecules, and therefore cannot easily recognize the similarity between a sidechain and a small molecule that share a functional group. We introduce a new method, called AtomPaint, that avoids these limitations by employing a fully-atomic representation of protein structure. Starting from a model of a desired binding interaction, our method proceeds by (i) converting that model to a 3D image, (ii) masking out the parts of that image that need to be redesigned, (iii) using a diffusion model to inpaint the masked voxels, then (iv) using a classification model to identify the amino acids in the inpainted image. Both models are SE(3)-equivariant ResNets, and were trained on a dataset of structures from the Protein Data Bank (PDB) curated to emphasize protein/non-protein interactions. In a sequence recovery benchmark, AtomPaint performed better than random guessing, suggesting that it understands some aspects of molecular structure. We discuss possible avenues of improvement, in the hopes that the advantages of our novel image-based approach can be fully realized.

## Introduction

The field of computational protein design has advanced to the point where we can reliably design proteins that adopt basic folds [1, 2, 3, 4]. Despite this, many kinds of protein functions remain difficult to design. This work focuses on one of these functions: binding to non-protein entities such as DNA, RNA, and small-molecules. An algorithm capable of reliably designing these binding interactions could help to build a variety of useful proteins, including enzymes, biosensors, transcription factors, binders for post-translationally modified proteins and gene editing tools.

Many of the machine learning algorithms used for protein design and protein structure prediction, including ProteinMPNN [5], RFdiffusion [3], and AlphaFold2 [6], can only be used for systems composed entirely of protein. The reason is that these methods use coarse-grained representations of protein structure where amino acids, rather than atoms, are the smallest building block. More specifically, these methods view proteins as graphs, where each node is an amino acid and each edge is some sort of spatial relationship. While this kind of representation cannot accommodate small molecules, it does have some important benefits. First, it makes it computationally tractable to work with systems that are big enough to be useful, i.e. hundreds of amino acids rather than hundreds of atoms. Second, in any context where mutations are being considered, the number of amino acids is known and fixed while the number of atoms is not.

Algorithms for designing protein/non-protein interactions must instead use representations capable of accommodating arbitrary molecules. The most common approach, used by LigandMPNN [7], RoseTTAFold-AA [8], Boltzdesign1 [9], AlphaFold3 [10], and others, is to use a heterogeneous graph where some nodes represent amino acids (or nucleotides) while others represent individual atoms. This approach maintains most of the benefits of a fully coarse-grained representation, but it still has drawbacks. The biggest relates to the fact that there are far more examples of amino acids than non-amino acids in the Protein Data Bank (PDB). To make efficient use of this data, then, a model would ideally be able to learn rules that generalize from amino acids to non-amino acids. For example, this might mean recognizing common structural motifs such as H-bonds, salt bridges, buried non-polar surface area, pi-stacking, and more. This might also mean recognizing functional groups that commonly occur in both small molecules and amino acids, such as alcohols, amines, amides, carboxylic acids, benzyls, phenols, and more. Such generalizations can only happen if amino acids and non-amino acids are represented in the same way, though. Another drawback is that heterogeneous representations have more pairwise relationships to learn. For example, with 20 amino acid, 8 nucleotides, and 6 atom types, there are 1156 possible relationships with different spatial characteristics, some of which may not occur frequently in the training data. In contrast, with 6 element types, there are only 36 possible relationships.

In this work, we present an new algorithm, called AtomPaint, for designing protein/non-protein interactions. The distinguishing feature of this algorithm is that it uses a full-atom representation for proteins and non-proteins alike, for the purpose of learning more realistic rules concerning protein/non-protein interactions. The key change enabling this feature is that we encode macromolecules as 3-dimensional (3D) images, not as heterogeneous graphs. This encoding allows the use of convolutional neural network (CNN) models, which are fast enough to process images containing thousands of atoms. We trained two models: one generative model capable of inpainting (i.e. filling in missing patches of) such images, and one classification model capable of identifying an amino acid within such an image given the position of its backbone atoms. To design a new macromolecular interfaces, we start with a rough model of the desired interface, erase the parts of the interface that we believe to be incorrect, fill in those erased parts using the generative model, then extract the protein sequence of the new interface using the classification model (Figure 1). This approach is unique in that it allows the model to learn its own coarse-grained representation, and predicts both conformation and sequence at the same time.

**Figure 1.**
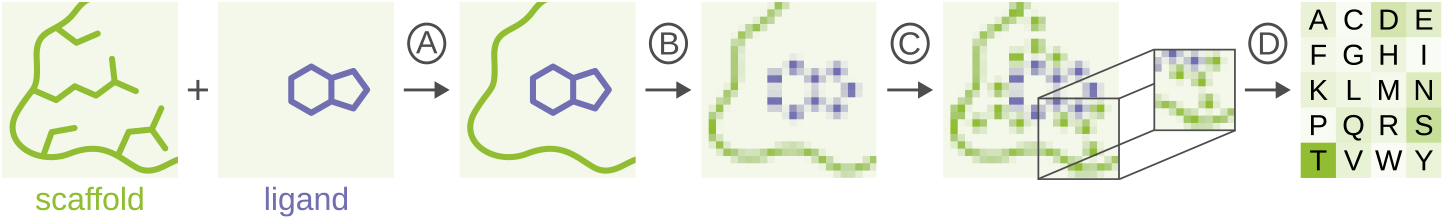
An overview of the design process. A) Input model creation: This could be done in different ways. Here, we illustrate the creation of a binding site for a ligand (purple) in an existing scaffold protein (green). Note that this model contains the native backbone, but not the native sidechains. B) Voxelization: The input model is converted into a 3D image. C) Inpainting: A diffusion model fills in any atoms that are missing from the input model. D) Amino acid classification: A small region around each unknown amino acid is cropped, and a classification model is used to identify the amino acid in each crop. In this example, threonine (T) is shown to be the most probable amino acid in the highlighted position. This step is repeated for each unknown amino acid in the image, and the result is a new protein sequence.

## Results

Let us begin by clearly defining the problem that AtomPaint is meant to solve. The goal is to modify an existing protein (which we will refer to as the scaffold) to bind a new non-protein partner. The required inputs are (i) a structural model of the desired binding interaction and (ii) a list of the positions that are allowed to mutate. The input model would typically be constructed by starting with a structure of the scaffold and manually replacing an existing binding partner with the new one (Figure 1a). In this way, the new binding partner would be positioned correctly relative to the scaffold, but would not yet be compatible with the existing interface. Given these inputs, AtomPaint will output a mutated sequence that is predicted to lead to a functional binding interaction.

The first step towards making this prediction is to convert the input model into an 3D image, with voxels that each represent 1 Å^3^ volumes of space and channels that represent different element types (e.g. carbon, nitrogen, oxygen, etc.) (Figure 1b). The whole image is 35 Å in each dimension, which is large enough to fit a protein of about 200 amino acids. The voxels are filled in proportion to their over-lap with the atoms in the input model. Next, we create a mask to cover the parts of the interface that need to be redesigned. This step could be done in different ways, but our usual approach is to mask any voxels that overlap with any reasonable sidechain conformation for any of the mutable sequence positions, and then to unmask any voxels near the backbone. Unmasking the backbone prevents it from moving during the design process. The amino acid classification step, explained below, currently requires a fixed backbone, but in principle a more sophisticated classifier could accommodate a flexible backbone.

Once we have an image of the binding site and a mask, we can use a diffusion model to fill in the masked voxels in a manner that is consistent with both the unmasked voxels and the training data (Figure 1c). This process is called inpainting. The result is a new image containing a redesigned binding interface. We will explain the diffusion model in more detail below, but to give a brief overview, it is trained to remove noise from images. It can then be used to generate new images by starting from white noise and iteratively removing noise until a clear image emerges. Inpainting proceeds similarly, except on each iteration the voxels in the unmasked region are reset to their known values, from the input image. In this way, only the masked region is filled in.

After the diffusion model generates an image of the new binding site, the last step is to determine the sequence of the protein in that image. We accomplish this using a model trained to classify amino acids in images generated by the diffusion model (Figure 1d). The input to this model is an 11 Å crop of the generated image, with an additional channel that indicates the location of the C_α_ for the amino acid of interest. The amino acid is not necessarily centered within the image, nor is it in any particular orientation. However, the crop is constructed in such a way as to have a >95% chance of containing all of the amino acid’s sidechain atoms. Note that this manner of making crops relies on knowing the backbone coordinates for each amino acid, which is why we unmask the backbone for the inpainting step. The output from the model is a probability for each of the 20 standard amino acid types. Because this model only classifies one amino acid at a time, it has to be invoked separately on each position that is allowed to mutate. Note that these invocations are completely independent of each other. The entire sequence (and structure) is constructed simultaneously by the diffusion model; the classification model only reveals the sequence that is already present in the resulting image.

### Dataset

To train the models introduced above, we curated a new dataset from the macromolecular structures in the PDB. We had two requirements for our dataset: it needed to be comprised of images, and it needed to emphasize protein/non-protein interactions. We could not find an existing dataset that met both criteria, which is why we curated our own. Our basic strategy was to divide each biological assembly in the PDB into 10 Å grid cells, and to keep only those cells that were high-quality and non-redundant (by the metrics summarized in the following paragraph, and described fully in the Methods section). To generate a training example from this dataset, we would sample a random cell, sample a random coordinate frame within that cell, then make an image centered on and aligned to that frame. These images could include atoms from outside their cells; the cells were only used to roughly indicate the regions of structures that make for good examples.

To decide which grid cells to include in the dataset, we began by assigning a quality ranking to each structure in the PDB. The primary metrics in this ranking were resolution (applicable to crystal-lography and electron microscopy structures) and clash score [11] (applicable to all structures), but we included enough secondary metrics to avoid any ties. We also assigned a quality ranking to each assembly within each structure, based mostly on the biological relevance annotations provided by the depositors. Next, we clustered every protein and nucleic acid using a threshold of 80% sequence identity with >80% coverage, and every small molecule by identity. We then iterated through each assembly in ranked order, overlaid a grid on the assembly, and included each grid cell that both (i) had a density between 40–70 atoms/nm^3^ within a 20 Å radius and (ii) was within 10 Å of either a cluster or a pair of clusters that hadn’t yet been included in the dataset. The cluster pair criterion is important. It allows for the same binding interface to be included multiple times, as long as each time is with a different binding partner. The final dataset comprised 203,053 grid cells from 34,278 structures. Finally, we divided the dataset into training, validation, and testing splits such that any two structures with an Interpro [12] domain or family in common must both go into the same split.

Having given this overview of our dataset, we can comment more on the goals we initially set out for it. The first was to produce images. This is different from the datasets used for graph-based models. For example, the individual data points making up the AlphaFold3 dataset are chains (or pairs of chains) that have been cropped to 384 tokens (where a token is either an amino acid, a nucleotide, or an atom) [10]. This kind of data is not suitable for our purposes because (i) an entire chain may not fit in the image, so we still need to decide where the image should go, (ii) parts of a chain that are more than 384 tokens apart may still be in close physical proximity, and (iii) there may be interfaces that involve more than two chains. Our grid cell approach solves all of these problems by using spatial coordinates, rather than chains, as the fundamental data type. Our second goal was to emphasize protein/non-protein interactions. Of course, the first step towards achieving this goal was to not filter out all non-protein entities, as many existing datasets do. Beyond that, the way we filtered the grid cells was explicitly designed to enrich for these kinds of interactions, without adding excessive redundancy to the dataset.

### Neighbor location pretext task

Because diffusion models are expensive to train and difficult to evaluate, we devised a simpler task to use for our initial experimentation with model architectures. This task is based on an idea put forth by Doersch et al. [13], namely that you can create a self-supervised task from an unlabeled image by making two nearby crops of that image, then having a model predict the location of one crop relative to the other. To make accurate predictions, the model must be able to extrapolate how the features visible in each crop extend outwards and intersect with each other. We applied this same idea, but in the context of 3D macromolecular images, and called it the neighbor location pretext task. Our crops were 15 Å large, and were separated by a random translation of 1–5 Å and a random rotation of up to 20°. The second crop could be in one of six locations relative to the first: above, below, left, right, in front, or behind (i.e. one location corresponding to each face of a cube). The model makes predictions by assigning a probability to each of these six relative locations.

There are two features that make this task good for experimenting with model architectures meant ultimately for a diffusion model. First is that this task is not protein-specific. A common protein-specific task is to predict the identity of an amino acid given the structure of its immediate environment (excluding its sidechain) [14, 15, 16, 17]. This task would be fast to train and easy to evaluate, but it wouldn’t necessarily reflect how a model architecture would perform on the task we truly care about, which is to design binding interfaces for *non-protein* partners. Second is that this task makes use of data from the entire PDB. This is the advantage that self-supervised tasks have over supervised ones. A common supervised macromolecular task is to predict the protein/small-molecule binding affinities collated in the PDBbind dataset [18] This would also be fast to train and easy to evaluate, and not fully protein-specific, but it is limited to a small number of training examples for which binding affinities have been experimentally measured (around 10,000). Many of these training examples are also quite redundant, featuring the same protein binding slightly different molecules.

Figure 2 summarizes the results achieved by several noteworthy model architectures on the neighbor location task. The first comparison to make is between the standard ResNet (orange) and the SE(3)-equivariant ResNet (red). Briefly, SE(3)-equivariance means that the model will recognize features regardless of what orientation they appear in, subject to some error due to way the image is voxelized [19]. In keeping with the fact that SE(3)-equivariant models have this additional inductive bias, they also have significantly fewer parameters than their non-equivariant counterparts. In principle, on systems like macromolecules that are not intrinsically changed by rotation, SE(3)-equivariant models should learn faster and generalize better. For more discussion on SE(3)-equivariance, particularly in the context of image-based models such as ResNets, refer to the supplemental material. Returning to the ResNets shown in Figure 2, we found that the equivariant implementation significantly outperformed the non-equivariant implementation. Both ResNets had the same number of layers, and the same latent variable dimensions at each layer. However, the equivariant ResNet had 24.7 times fewer parameters (713K versus 27.6M).

**Figure 2.**
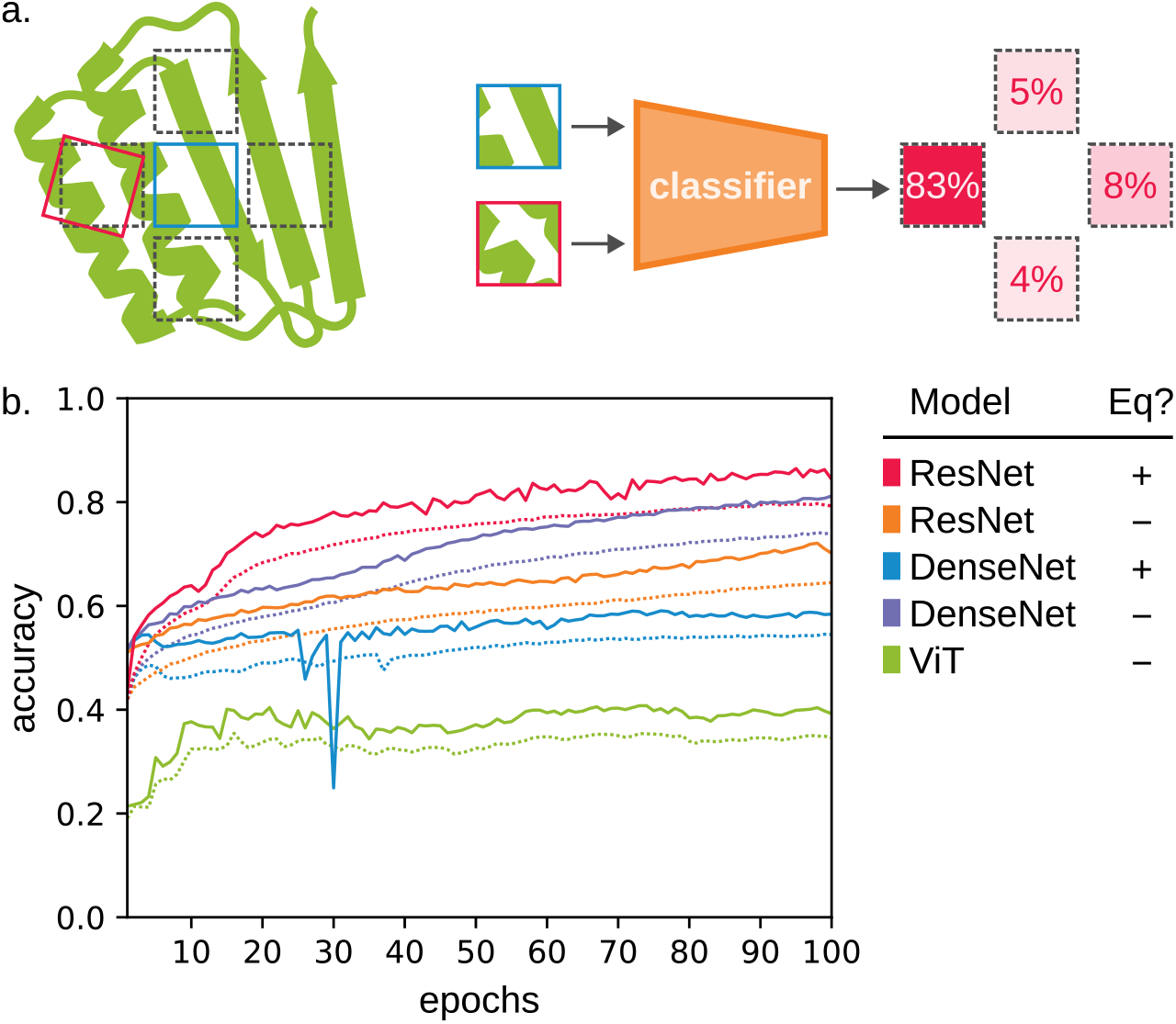
Using a pretext task to find reasonable model architectures. A) An illustration of the neighbor location task. The green cartoon represents a macromolecule. The blue and red boxes represent the first and second crops, respectively. The black dashed boxes represent the possible locations for the second crop. For clarity, this illustration shows only 4 possible locations, but the actual task has 6. Notice that the second crop is offset from its corresponding location by a small random translation and rotation. The only inputs to the classifier are the two crops. The outputs are probabilities for each location. In this illustration, an 83% probability is assigned to the correct location. B) Training curves for select model architectures on the neighbor location task. Solid and dashed lines represent the validation and training sets, respectively. The “Eq?” column in the legend indicates whether or not a model is SE(3) equivariant.

The next comparisons to make are between the ResNets (red, orange) and the DenseNets (blue, purple). Both ResNets and DenseNets are established image-processing model architectures that involve “skip” connections—connections between non-adjacent layers that are believed to help gradients propagate through the entire model more easily. The primary difference is that ResNets use addition for the skip connections, while DenseNets use concatenation. Unlike with the ResNets, we found that the non-equivariant DenseNet outperformed the equivariant DenseNet. Both models had similar numbers of parameters (23.1M and 673K, respectively) and latent variable dimensions as their ResNet counterparts. The equivariant DenseNet exhibited more instability during training (see the temporary drop in validation accuracy around the 30_th_ epoch), which might partially explain the impaired performance. Regardless, we found that the equivariant ResNet architecture outperformed both DenseNet architectures, in this context.

The final model we will comment on here is the Vision Transformer (ViT, green). Briefly, ViTs work by breaking up an image into patches, then treating those patches in the same way that large language models (LLMs) treat words [20]. ViTs generally outperform convolutional neural networks (CNNs), including ResNets, on 2D image recognition tasks [21]. However, they require larger datasets and longer training times due to their weaker inductive bias. Specifically, they have neither rotational nor translational equivariance. Our dataset is very small by ViT standards, which may be why the ViT does not perform well. Our best performing ViT (shown here) has 9.5M parameters, 13.3 times more that the equivariant ResNet.

In summary, we developed a self-supervised task and used it to experiment with a variety of established model architectures. We also used it to experiment with hyperparameters including atom radius (Figure S1), element type (Figure S2), model width (Figure S5), block architecture (Figure S6), and the choice of equivariant nonlinearities (Figure S3, S4). From this, we concluded that an SE(3)-equivariant ResNet was the most promising model architecture.

### Diffusion model

Our next step was to develop and train a diffusion model capable of generating new images of macromolecular structures. As we mentioned briefly when introducing our neighbor location task, one of the challenges in training a diffusion model is finding a way to quantify how realistic its output is. For 2D images, a metric called Fréchet inception distance (FID) is commonly used for this purpose [22]. Briefly, FID compares how a pretrained classification model (canonically Inception v3 [23]) encodes a large set of reference images, to how it encodes a large set of generated images. The specific comparisons are between the distributions of the activations of the neurons in the last hidden layer of the model. A distance of zero indicates that the classifier cannot distinguish a generated image from a reference image. To implement this metric in the context of 3D macromolecular images, we used our best-performing model on the neighbor location task. Recall that this model takes as input two 15 Å crops separated by 1–5 Å and that the diffusion model generates 35 Å images. This means that we can sample 4 pairs of non-overlapping crops from each generated image. We evaluated this metric by adding normally-distributed noise to the position of each atom in a structure, and found that after 1000 samples (250 images), a distance of around 35 corresponded to standard deviation of 0.6 Å (Figure S7).

The architecture of our diffusion model is illustrated in Figure 3a. The overall structure is that of a U-Net [24]. The encoder (i.e. the first half) is very similar to the SE(3)-equivariant ResNet architecture that performed the best on the neighbor location task, except that it is deeper to accommodate the larger input image size. The decoder (i.e. the second half) is composed of regular ResNet blocks, without SE(3)-equivariance. The deepest of these blocks includes a self-attention layer.

**Figure 3.**
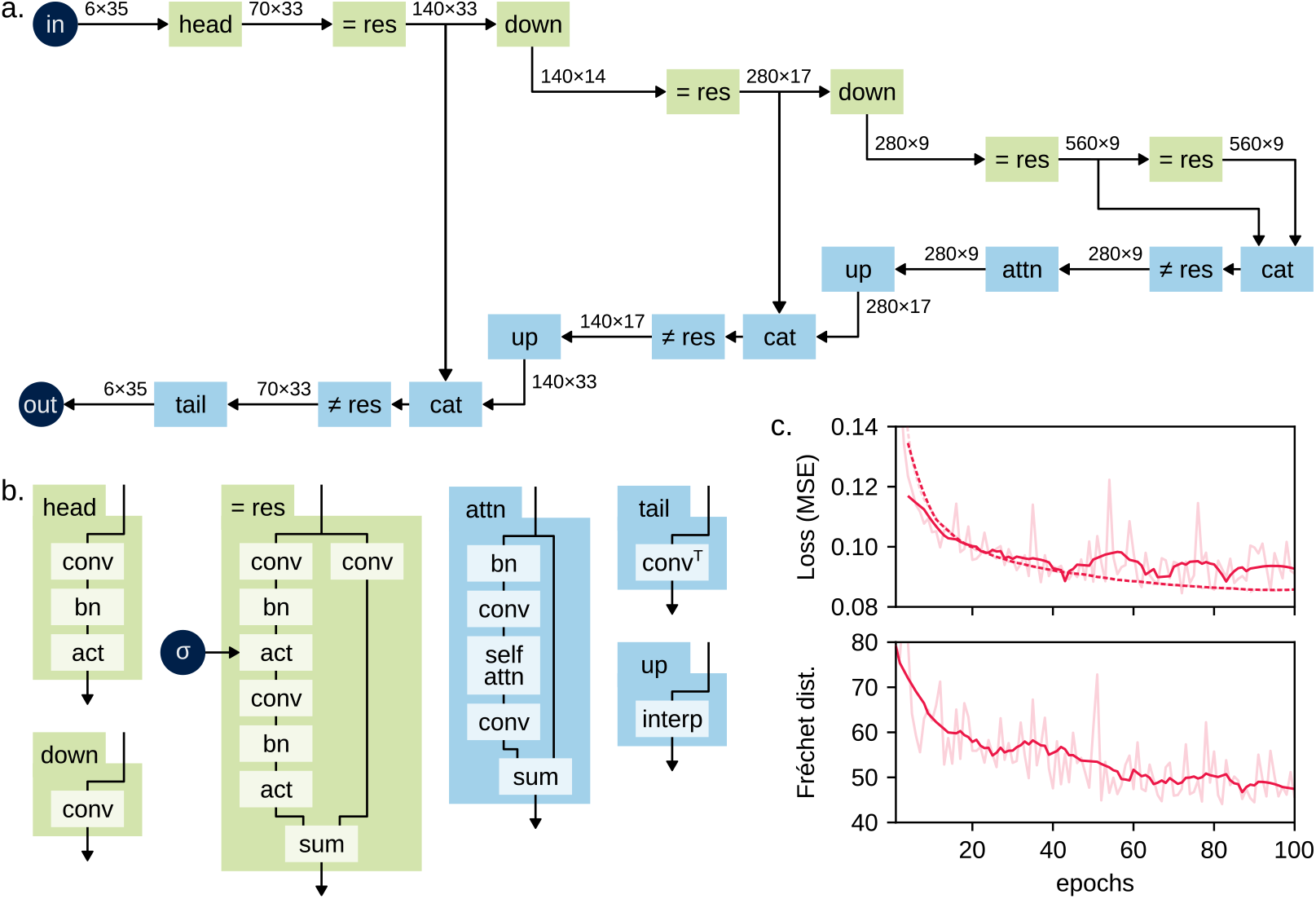
Training the diffusion model. A) A schematic of the U-Net model architecture. The input and output images are shown as navy circles. The blocks making up the model are shown as rectangles. The green blocks maintain SE(3) equivariance; the blue blocks do not. The “head” and “tail” blocks are the first and last blocks of the model, respectively. The “= res” and “≠ res” blocks are equivariant and non-equivariant residual blocks, respectively. The “down” and “up” blocks change the size of the spatial dimensions of their inputs. The “cat” blocks concatenate their inputs along the channel dimension. The “attn” block contains a self-attention layer. The arrows show how the blocks are connected. The shape of the latent representation is indicated above each arrow as “channels × voxels per spatial dimension”. Note that all of the latent representations have three equally-sized spatial dimensions, so the actual shape of—for example—the input is 6×35×35×35. B) More detailed schematics of the blocks making up the diffusion model. Here, “conv” is a 3D convolution, “conv ⍰” is a 3D transpose convolution, “bn” is a batch normalization, “act” is a nonlinear activation function, “sum” is simply a sum of two inputs, “self attn” is self-attention, and “interp” is a 3D linear interpolation. The green background denotes equivariant versions of these operations; the blue background denotes standard, non-equivariant versions. These schematics still elide some details; refer to the Methods section for a complete description of each blocks. The equivariant residual block has an additional input, which is the amount of noise present in the input image. We left this input out of the schematic in panel (A) for clarity, but it is present for every such block. The schematic for the non-equivariant residual block is not shown here. It is largely the same as that of the equivariant residual block, including the noise input, but there are some minor differences. Refer to the Methods section for details. C) Quality metrics plotted over the course of a training run. The upper plot shows mean-square error (MSE) loss on the training (dashed line) and validation (dashed lines) datasets. The lower plot shows the Fréchet distance metric. In both plots, the dark lines are smoothed, the faint lines are not.

We based our diffusion algorithm on the ideas of Karras et al. [25]. Some distinguishing features of this algorithm are (i) a choice of noise schedule that lends itself to flatter diffusion trajectories, (ii) a second order integrator that allows for larger diffusion steps, and (iii) a formulation that varies whether the model predicts the noise present in the input or a denoised version of the input, depending on the noise level.

In training our diffusion model, we found that our biggest challenge was to prevent overfitting. We addressed this in the following ways. First, we used weight decay, specifically for the weights (but not the biases) of the convolutions (Figure S9). This favors weights that are closer to zero and has a regularizing effect. Second, we used reduced learning rates (Figure S10). This improves training stability. Third, as shown in Figure 3, we used SE(3)-equivariant layers in the first half or the model (i.e. the encoder). The equivariant layers have fewer parameters, and without them we observe overfitting (Figure S8). Combining all of these strategies, we were able to train a model that achieved a Fréchet distance of 45 without overfitting (Figure 3).

It’s worth mentioning a few common diffusion modeling tricks that we tried, but did not observe any benefit from. First was exponential moving averaging (EMA). Specifically, we used an algorithm called Switch EMA, which accumulates averaged weights outside of the model over the course of an epoch, then substitutes those weights back into the model before the start of the next epoch [26]. This did have a regularizing effect, but also had much worse training loss, validation loss, and Fréchet distance (Figure S11).

The second and third tricks we tried were conditioning and self-conditioning. The idea behind conditioning is to provide an additional input describing the content of the image being denoised, which the model can use to make better predictions. We provided (i) the fractions of atoms that are protein, DNA, and RNA and (ii) the fractions of atoms belonging each of the 11 CATH architectures with more than 7500 members [27]. Self-conditioning is a similar idea, but the additional input is a denoised version of the noisy input image, generated by running the model itself [28]. During training, this doubles the number of times that the model needs to be invoked. During inference, this costs nothing, since the diffusion process is iterative and the output from the previous step is always available. Neither conditioning technique had a significant effect on training or validation loss, but both had a significant negative effect on Fréchet distance (Figure S12).

The fourth trick we tried was to (i) compress the input image into a smaller latent space, (ii) perform diffusion in that space, then (iii) decompress the result to get the output image [29]. The reason for doing this is to make the diffusion model smaller, leading to faster training and inference times. We trained variational autoencoders (VAE) to encode and decode the input images, but we were not able to get good reconstruction performance (quantified using mean squared error, MSE) for those models with significant compression ratios (Figure S13). Anecdotally, a common pathology was for 70-80% of the image to be reconstructed well, but for the remaining voxels to be filled in as a solid block.

### Amino acid classification model

The final step in our design protocol is to identify the amino acids in the images produced by the diffusion model. As mentioned above, we accomplished this by training a model to identify an amino acid, given a crop of such an image and the location of that amino acid’s C^α^. Figure 4 shows both the architecture and the performance of the best model we found. This model is SE(3)-invariant, meaning that it produces the same output regardless of how the input is rotated. Note however that we found many model architectures that achieve >95% validation accuracy (with balanced classes) on this task (Figure S14) in less than one full epoch. This likely reflects that it is not especially difficult to identify an amino acid given its structure; humans can perform this task with 100% accuracy.

**Figure 4.**
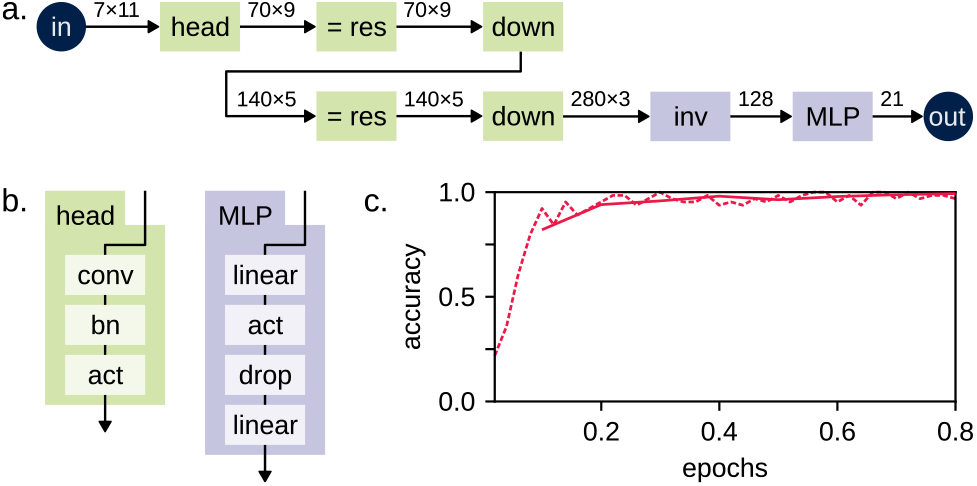
Training the amino acid classifier. A) A schematic of the classification model. The inputs and outputs are shown as circles, and the blocks making up to model are shown as rectangles. The green blocks are SE(3)-equivariant, and the purple blocks are SE(3)-invariant. The “down” block reduces the size of the latent representation. The “= res” block is identical to that in Figure 3. The “inv” block converts an SE(3)-equivariant image into an SE(3)-invariant vector. Refer to the Methods section for more details. The “MLP” block is a multi-layer perceptron that calculates probabilities for each of the possible output classes. The numbers above the arrows give the shape of the latent representation at each step, as in Figure 3. B) More detailed schematics of the blocks making up the classification model. “conv”, “bn”, and “act” are all defined as in Figure 3. “linear” is a fully-connected linear layer, and “drop” is a dropout layer. C) Improvement in the model’s accuracy over the course of a training run. The solid and dashed lines represent the validation and training datasets, respectively. For both datasets, the amino acids are weighted such that each amino acid is equally likely to occur.

Despite this not being a difficult task, we found that several aspects of data preparation were essential for good performance. First, the crops must not be too big. We observed the best results with 11 Å crops (Figure S15). Second, the crops must contain most of the sidechain atoms being classified (Figure S16). This latter determination must be made using only the positions of the backbone atoms, as the positions of the sidechain atoms are not known at inference time. We approached this by finding, in a coordinate frame defined by the backbone atoms, the 4 Å radius sphere that contained the greatest fraction of sidechain atoms in a sample of residues drawn from the PDB. This sphere was located roughly in the direction of the C_β_ atom, but 3.3 Å from the C_α_. We then required that this entire sphere (diameter: 8 Å) be contained within the cropped image (side length: 11 Å).

### End-to-end training

Having independently trained both a diffusion model and an amino acid classification model, we sought to improve both models by training them end-to-end. This means that during training, the output of the diffusion model was used as input to the amino acid classifier, and then the weights for both models were updated using the loss functions for both tasks. As such, the diffusion model learns to output images that can be recognized by the classifier, and the classifier learns to accommodate the kinds of errors made by the diffusion model. To balance the contributions of the two loss functions, we scaled each using a learnable “uncertainty” parameter [30]. During end-to-end training, we observed that both loss functions improved simultaneously and without overfitting (Figure 3a,b).

We evaluated the complete AtomPaint design protocol by measuring sequence recovery [5], a standard metric for fixed-backbone protein design algorithms. This metric works by starting with a known structure, removing all of its sidechain atoms, then predicting the identities of its amino acids. The reported value is the fraction of amino acids that are correctly predicted, after balancing the frequency with which each amino acid appears. We calculated sequence recovery by converting the initial structure into an image, creating a mask that covers the sidechains but not the backbone, using our diffusion model to fill in the masked voxels, then using the amino acid classification model to predict amino acid identities. Unfortunately, AtomPaint does not perform well by this metric. When the dataset is weighted such that each of the 20 amino acids are equally probable, AtomPaint only makes the correct prediction 5.7% of the time. For reference, a uniformly random guess would be correct 5% of the time. The primary problem is that 83.4% of AtomPaint’s predictions are glycine (Table S1). Upon visual inspection, we found that these predictions occur because the diffusion model fails to generate any sidechain atoms close enough to the C_α_, not because the classifier fails to identify the sidechains generated by the diffusion model.

Despite the poor sequence recovery, there are signs that the model has truly learned some aspects of macromolecular structure. First, if we compare AtomPaint to a model that picks amino acids randomly, but with the same frequencies as AtomPaint, AtomPaint is correct more often for every amino acid except cysteine (Table S1). Second, we anecdotally observe that AtomPaint often correctly prefers nonpolar atoms in protein cores and polar atoms on protein surfaces. Figure 5c shows a representative example of this. Note the preponderance of carbon voxels (green) above and below the buried β-sheet, and the preponderance of nitrogen (blue) and oxygen (red) voxels on the surface. Third, AtomPaint is capable of generating recognizable sidechains. Several examples of this are shown in Figure 5d-g. These examples are all from cases where the predicted amino acid matches the actual amino acid, but of course there are also cases where AtomPaint confidently predicts the wrong amino acid.

**Figure 5.**
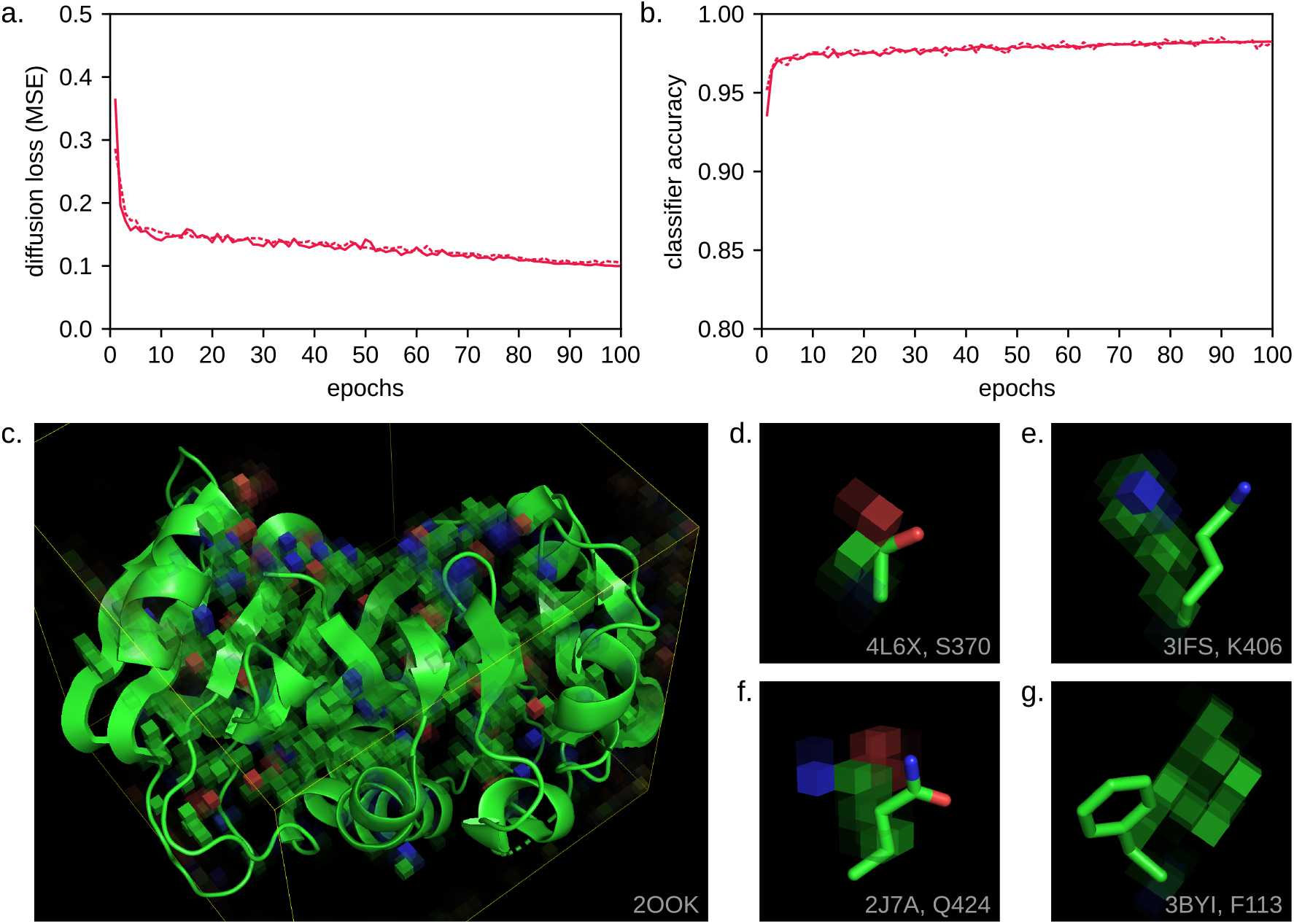
Training the end-to-end model. A) Image reconstruction loss achieved by the diffusion model during training, measured on the training (dashed line) and validation (solid line) datasets. B) Accuracy achieved by the classification model during training, measured on the training (dashed line) and validation (solid line) datasets. C) Inpainting results for a representative protein. In this example, the backbone atoms were given and the sidechain atoms were not. Only the voxels generated by the diffusion model, i.e. those representing sidechains, are shown. The green, red, and blue voxels represent carbon, oxygen, and nitrogen atoms, respectively. The yellow lines delineate the boundaries of the 35 Å image. The protein is shown as a cartoon. D-G) Close-up views of sidechain conformations predicted by the end-to-end model. Voxel colors are the same as in panel C. The true sidechain conformations are shown as sticks.

## Discussion

In this work, we introduce AtomPaint, an approach to designing protein/non-protein interactions that— to our knowledge—has never been published before: make an image of the desired interaction, erase any voxels that may not be correct, use a generative model to fill in those voxels, then extract the sequence of the protein in the resulting image. The motivating advantage of this approach is that the same full-atom representation is used for both protein and non-protein entities, which allows the model to learn rules that generalize to both. However, this approach has disadvantages as well. First, for protein-only design problems, it is more complicated than necessary. Second, it assumes that a desired binding geometry is known. Not all design tasks satisfy this assumption, but many do. A typical example might be adapting an enzyme to bind a new substrate, particularly if the old and new substrates can be aligned by some shared chemical motifs. Third, this approach cannot consider alternate conformations of non-protein entities. In theory, this restriction could be lifted by training a model to recognize whether an image contained a particular small-molecule, given a topological description of that molecule (e.g. a SMILES string). Such a model could then be used to condition the diffusion model to generate images that contain any molecule of interest. In practice, though, getting this scheme to work well would not be a trivial exercise.

We have described our efforts to optimize AtomPaint, but many avenues for improvement remain. We will comment on several of these avenues in the paragraphs to follow, beginning here with the dataset. The most significant problem we faced while training the diffusion model was overfitting. This ultimately limited the size of the model we could train. The ideal way to address overfitting is to make the training data larger and more varied. Fortunately, there are ways to do this. First, instead of clustering structures and keeping only the highest quality representative of each cluster, as we did, one could weight the members of each cluster by quality. This would ensure that many more structures appear in the training data, while still maintaining a preference for high-quality structures. Second, we could incorporate computed structure models (e.g. AlphaFold3 models) into the dataset. We did not take this approach initially because, prior to the release of AlphaFold3, such models were protein-only. Since our main goal was to design protein/non-protein interfaces, we did not want to dilute the already limited number of structures in our dataset that featured non-protein entities. Now, however, computed structure models can include all kinds of molecules, and could be a valuable source of data.

Another avenue for improvement is the diffusion model itself. We observed the best results with partially SE(3)-equivariant models. That is, models where the encoder maintains equivariance and the decoder does not. However, we still believe that fully equivariant models have the potential to produce the best results. One possible explanation for why these models have not lived up to their potential is that the equivariant nonlinearities available at present seem to be much less expressive than standard nonlinearities like the rectified linear unit (ReLU). This is illustrated by Figure S3, which shows the mostly linear relationship between inputs and outputs for a number of equivariant nonlinear functions. If improved equivariant nonlinearities are developed in the future, they might enable fully equivariant models to outperform partially equivariant ones. We did not extensively experiment with altering the ratio of equivariant to non-equivariant layers. It is possible that this could be an important hyperparameter, as well.

The last avenue for improvement that we’ll discuss here is the amino acid classification model. Despite performing with near-perfect accuracy on data from the PDB, several possible improvements to the overall pipeline involve this model. Before commenting on those improvements, though, it’s worth commenting on the task that this model is meant to solve: identifying an amino acid given an image and the location of a C_α_atom. This is similar to a task that has been extensively studied: identifying an amino acid given an image and the location of a C_α_ atom, but with all of the amino acid’s sidechain atoms withheld from the image [14, 31, 15, 16, 17]. We will refer to these as the “sidechain-aware” and “sidechain-blind” tasks, respectively. The sidechain-blind task is more difficult, of course, but it’s still informative to compare some general features of the models trained on both tasks. First, sidechain-blind models typically require that the image be centered on and aligned to the amino acid in question. Due to the way our images are generated (i.e. by a diffusion model), our sidechain-aware model must be able to accommodate amino acids in any orientation, and that aren’t perfectly centered. Second, sidechain-blind models typically accept 20 Å inputs, with 1 Å voxels. We found that smaller 11 Å inputs, still with 1 Å voxels, worked better for our use case, likely because less context is required when the sidechain itself is given. Third, our sidechain-aware model is SE(3)-invariant. Most of the sidechain-blind models are not. However, Weiler et al. [31] show that SE(3)-invariance leads to improved performance on this task.

Let us return to our discussion of avenues for improvement involving the amino acid classification model. The first is to predict sidechain torsion angles in addition to amino acid identities. In an end-to-end training run, requiring the classification model to have a more detailed understanding of the structure might encourage the diffusion model to generate more realistic structures. The second improvement is to allow flexible backbone design. Currently, the diffusion model is capable of generating images with new backbone conformations, but the classification model is not yet capable of extracting protein sequences from images with unknown amino acid positions. To relax this requirement, the classification model must gain the ability to predict both the position and the order of the amino acids to identify. This could be implemented as a Markov process. That is, the model could initially be given the position of the last known, fixed-backbone amino acid and asked to predict the position of the first unknown, flexible-backbone one. This information could then be used to locate the next amino acid, and so on. In the same manner as the side chain torsion angles, training the classification model to make these predictions might also help encourage the diffusion model to make better images.

In conclusion, we have presented a new method for computational protein design that emphasizes protein/non-protein interactions. Compared to existing machine learning methods, ours is unique in that represents both protein and non-protein atoms in the same way. This allows the underlying neural networks to learn their own coarse-grained representations of macromolecular structure, and to find general rules that apply to both proteins and non-proteins, ameliorating the fact that the former are more common that the latter in the training data. The version presented here is for fixed-backbone design with known binding site geometries, but the same underlying generative model could accommodate both flexible backbones and mobile ligands. We believe that the results shown here establish image-based diffusion models as a promising method for protein design, deserving of further exploration.

## Methods

### Source code

Source code for all of the software developed for this project is publicly available:

- https://doi.org/10.6084/m9.figshare.30842459
- https://doi.org/10.6084/m9.figshare.30826517
- https://github.com/kalekundert/atompaint
- https://github.com/kalekundert/macromol_census
- https://github.com/kalekundert/macromol_gym
- https://github.com/kalekundert/macromol_voxelize

### Dataset

In this section we use the terms “structure”, “model”, “biological assembly”, and “entity” in the same manner as they are used in the PDBx/mmCIF specification. We use the terms “chain” and “subchain” to refer to what the specification calls `auth_asym_id` and `label_asym_id`, respectively.

We downloaded a copy of the PDB, along with validation reports, on December 4, 2024. We then ranked the quality of each structure according to the following criteria, in order: (i) resolution, rounded to the nearest 0.1 Å, for crystal and electron microscopy (EM) structures with resolution better than 4 Å; (ii) clash score, rounded to the nearest 0.2, for all structures; (iii) the number of NMR restraints, for NMR structures; (iv) *R*_free_, for crystal structures; (v) Q-score, for EM structures; (vi) deposition date, with later being preferred; (vii) PDB identifier. These criteria ensure a deterministic ranking for every structure in the PDB. From each structure, we then extracted biological assemblies according to the following criteria: (i) biological relevance, i.e. assemblies of type “representative helical assembly”, “complete point assembly”, “complete icosahedral assembly”, “software defined assembly”, “author defined assembly”, or “author and software defined assembly”; (ii) complete coverage of all subchains in the structure, (iii) size, (iv) order of appearance. Collectively, these steps yielded a deterministically ranked list of every biological assembly in the PDB.

We then clustered every entity in the PDB. We clustered proteins and nucleic acids using MMseqs2 [32] with a threshold of 80% sequence identity and 80% coverage (coverage mode 0, cluster mode 0). These thresholds were determined by manually inspecting clusters, with the goal of finding the lowest threshold that would still only cluster entities with near identical structures. We clustered every other entity by identity. For branched entities, we considered two entities identical if their molecular graphs were isometric.

We manually classified the 802 most common small molecules in the PDB as either “specific”, “non-specific”, or “non-biological”. “Specific” are those that bind to a larger molecule (typically a protein) at a defined site, in a defined orientation. “Non-specific” are those that bind, possibly in a defined region, but not in a defined orientation. Many of the molecules to get this classification were lipids. “Non-biological” are those that are not present in nature, but were added to assist with determining the structure. Some small molecules belong to different categories in different structures. For example, glucose is bound specifically by many lectins, but is also used as a crystallization aid. A lipid that is part of a non-specific membrane in one structure might be bound specifically to a flipase in another. In such cases, we assigned the label that seemed to best describe the greatest number of structures. We then automatically classified another 2210 small molecules, for a total of 3012, by clustering every small molecule in the PDB such that any two molecules with >90% Tanimoto similarity would belong to the same cluster. If a cluster contained one manually classified molecule, then every molecule in that cluster was assigned that label. If a cluster contained multiple manually classified molecules, then the least “specific” of those labels was used.

Next, we created our training examples. We began by iterating through each biological assembly in ranked order. We skipped any assemblies that (i) contained any elements other than C, N, O, S, Se, P, Mg, Ca, Mn, Fe, Co, Ni, Cu, Zn, or (ii) exceed a density of 70 atoms nm^−3^ in any location, as this is often indicative of erroneous structures with overlapping subchains. We also skipped 8h2i and 5zz8. Both of these structures have biological assemblies that erroneously leave large amounts of empty space between each asymmetric unit, and as a result required prohibitive amounts of memory to process. On each remaining assembly, we superimposed a 10 Å grid. We aligned the grid to the first three principle components of the coordinates of every atom in the assembly. We then iterated through each grid cell with density greater than 40 atoms nm^−3^ and flagged those that contained a unique entity (i.e. an entity whose cluster is not yet present in the dataset) that either comprises at least 75% of the atoms in the cell (i.e. the entity is larger than the cell, but occupies most of it), or has more than 75% of its atoms within the cell (i.e. the entity is smaller than the cell, but mostly contained within it). We also flagged cells that contained a unique pair of entities, both of which individually either comprise at least 25% of the atoms in the cell, or have more than 75% of their atoms within it. After processing all of the grid cells for one biological assembly, we added the flagged grid cells to the dataset. Our final dataset comprised 34,278 structures, 34,583 biological assemblies, and 203,053 grid cells.

To create training, validation, and test splits, we queried InterPro for the set of domains and families found in each structure. We clustered every structure in our dataset such that any two structures with at least one domain or family in common would be part of the same cluster. We then randomly assigned each cluster to one of the splits, with probability proportional to difference between the target number of grid cells in each split (80% training, 10% validation, 10% test), and the number of grid cells that would be in that split if the current cluster were assigned to it. The result was a training split with 26,496 structures and 162,442 grid cells, a validation split with 4,023 structures and 20,305 grid cells, and a test split with 3,759 structures and 20,306 grid cells. Throughout this work, we consider one epoch to be one pass through all of these grid cells.

### Neighbor location pretext task

The input to the neighbor location task was a pair of 3D images of adjacent regions from one biological assembly. Both images had the same shape. Both had 6 channels, each corresponding to different element types: C, N, O, P, S/Se, Mg/Ca/Mn/Fe/Co/Ni/Cu/Zn. Both were 15 Å on each side, with 1 Å voxels. To create the first image, we uniformly sampled a grid cell (i.e. a volume within a specific biological assembly that meets various quality criteria) from the appropriate split of our dataset. We then uniformly sampled a coordinate from within the grid cell to be the center of the image, and uniformly sampled a vector from the surface of a unit sphere to be the orientation of the image. For each atom within the image, we constructed a sphere centered on that atom with radius 0.5 Å. The value encoded by each voxel was the sum of the fractions of the volume of each sphere that intersected that voxel [33], for all spheres corresponding to atoms of the appropriate element type for the channel that the voxel belongs to. Note that the sum of all the voxels in the image would roughly equal the number of atoms in the image.

To create the second image, we first uniformly sampled one of the six directions corresponding to the cube faces of the first image. This direction is what the model would ultimately be trained to predict. We uniformly sampled a distance between 1–5 Å and placed the edge of the second image at this distance in the chosen direction from the edge of the first image. We then uniformly sampled an axis of rotation from the surface of a unit sphere and an angle between 0-20°, and rotated the second image by this angle around this axis relative to orientation of the first image. Finally, we filled in the voxels of the second image in the same manner as the first.

In each epoch, we chose a different random seed for all of the sampling steps required to make both images. This means that, while each epoch would feature images of the same general regions of the same macromolecules, the same exact images would not be reused.

We trained a number of different models on this task. All of the models we trained were composed of the same two components: an encoder and a classifier. The encoder would take an image as input and produce a flattened latent representation of that image (i.e. with spatial dimensions removed) as output. Most of our encoders were 3-5 layer CNNs that produced roughly 512-dimensional outputs, although this varied greatly. Each of the two input images would be fed independently into the same encoder (i.e. with the same weights, and with gradients acculumating) to produce two latent representations, which would subsequently be concatenated. The classifier would take the concatenated 1D latent representations as input and produce six logits, one corresponding to each of the possible directions relating the first and second images, as output. Typically, our classifiers were multilayer perceptrons with roughly 1024-dimensional input, one 512-dimensional hidden layer, and 6-dimensional output. Each linear layer of the MLP (except the last) was followed by a rectified linear unit (ReLU) nonlinearity and a dropout layer with a drop probability of 20%. We used cross entropy loss, and the Adam optimizer with a learning rate of 0.001.

### Diffusion modeling

We trained our diffusion models on 3D images of regions of macromolecular structures. We created these images in the same way that we created the “first” image from the neighbor location task, as described above, with two differences. First, the images measured 35 Å in each dimension. The voxels were still 1 Å, so these images simply had more voxels. Second, these images were scaled by a factor of 20. Empirically, this factor brought the standard deviation of all the voxels in all of the images in the dataset to approximately 1.

We used a training regimen based on the ideas of the EDM framework [25]. Briefly, this means that for each noise-free input image, we sampled a noise level from Σ *∼ e*^*N*(*μ*=−1.2, *σ*=1.2)^, where *N*(*μ, σ*) is the Gaussian distribution with mean *μ* and standard deviation *σ*. We then created a noisy image by adding noise sampled from *N*(0, Σ) to each voxel of the noise-free image. This noisy image, along with the noise level itself, was the input to the diffusion model. The output from the model was another image of the same shape as the input. To get the predicted noise-free image, we calculated a linear combination of the noisy input image and the output of the diffusion model. The coefficients of this linear combination were a function of the noise level, such that the model was mostly responsible for predicting the noise-free image when the input had significant noise, and mostly responsible for predicting the noise itself when the input had little noise. Our loss function was the mean squared error between the true noise-free image and the predicted noise-free image. For more details, refer to the above citation.

We used an algorithm based on both EDM [25] and RePaint [34] to generate images using our trained diffusion models. For generating unconditioned images (i.e. without inpainting), our algorithm was equivalent to the EDM algorithm, but with parameters optimized for our application. We used 98 diffusion steps, with an initial noise level of *σ*_max_ = 65.765, a final noise level of *σ*_min_ = 0.0279, and the noise schedule determined by the parameter *ρ* = 7.539. We added churn for all noise levels, with *S*_churn_ = 12.393. We performed two model evaluations in every step, as prescribed by Huen’s second-order method for solving differential equations. For generating conditioned images (i.e. with inpainting), we modified the EDM algorithm to incorporate ideas from the RePaint algorithm. Specifically, we applied the mask at the beginning of each diffusion step, and repeated the diffusion step for each noise level 10 times.

### Fréchet distance metric

To make a Fréchet distance metric, we began by training a classification model on the neighbor location task. As described in the above section on that task, this model was composed of an encoder and a classifier. The encoder was an SE(3)-equivariant ResNet. To maintain SE(3)-equivariance in each layer of the ResNet, we used the convolution, batch normalization, and nonlinearity implementations provided by the ESCNN library [19], in lieu of the standard implementations. The input was a 3D image with 6 channels and 15 voxels in each spatial dimension. The encoder was composed of one head block, four residual blocks, and one tail block.

The head block was composed of a convolution, a batch normalization, and a Fourier nonlinearity. The convolution had a 3x3x3 kernel, a stride of 1, and a padding of 0. The output of the convolution had the representation *R*(2), which corresponds to 70 channels, where 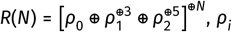 an irreducible representation of SE(3) with frequency *i*, ⊕ signifies a direct sum, and *ρ*^⊕*M*^ indicates that the representation *ρ* is direct-summed with itself *M* times. The Fourier nonlinearity was evaluated on a grid of 96 elements of SE(3), distributed evenly (i.e. by a numerical algorithm meant to maximize separation) and with octahedral symmetry [19]. This same grid was used for all of the Fourier nonlinearities in the model. At each grid point, in the spatial domain, the exponential linear activation (ELU) was applied.

Each of the four residual blocks was composed of an optional downsampling layer, followed by two branches, followed by a residual connection. We will refer to the input and output representations of each block as *R*(*a*) and *R*(*b*). The downsampling layer was present in the second and fourth blocks, and was a convolution with a 3x3x3 kernel, a stride of 2, a padding of 1, and an output representation of *R*(*a*). The first branch was composed of a convolution, batch normalization, tensor product nonlinearity, convolution, and batch normalization. Both convolutions had 3x3x3 kernels, strides of 1, paddings of 1, and no bias. The output representation of the first convolution was *R*(*a*), while that of the second was *R*(*b*). The second branch was composed of a single convolution, with a 1x1x1 kernel, a stride of 1, a padding of 0, and an output representation of *R*(*b*). The residual connection was the sum of the outputs from both branches, which have the same shape and representation, followed by a Fourier nonlinearity with the first Hermite function 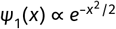 as the activation function. The four residual blocks had output representations *R*(3), *R*(5), *R*(7), and *R*(11), with the output from one block being the input to the next. Respectively, these representations had 105, 175, 245, and 385 channels.

The tail block, like the head block, was composed of a convolution, a batch normalization, and a Fourier nonlinearity. This convolution had a 4x4x4 kernel, a padding of 0, and an output representation of *R*(16), which corresponds to 560 channels. The stride doesn’t matter, because the input to this convolution is also 4x4x4. The batch normalization and Fourier nonlinearity layers are the same as in the head block.

The classifier was a multilayer perceptron (MLP) composed of a first linear layer, a ReLU nonlinearity, a dropout layer, and a second linear layer. The first linear layer had 560 × 2 = 1120 input channels and 512 output channels. The second linear layer had 512 input channels and 6 output channels. The dropout layer had a drop probability of 20%.

We calculated Fréchet distance using the 512-dimensional latent vector produced by the first linear layer of the classifier. We accumulated reference statistics for these vectors—specifically the running mean, the covariance times the number of observations, and the number of observations—over the entire training split of our dataset. Note that these statistics both contain the information necessary to calculate Fréchet distance and avoid various instabilities in the floating point arithmetic required to do so. To evaluate a diffusion model during training, we used the diffusion model to generate 384 35x35x35 images at the end of each epoch. We split each image into 4 non-overlapping pairs of 15x15x15 images, with each image separated from its partner by 3 voxels and from the edge of the larger image by 1 voxel. We then used this pair of images to produce a latent vector. With 384 images and 4 latent vectors per image, we produced 1536 latent vectors per epoch. We accumulated the statistics over these vectors in the same manner as we did for the training set, then reported the Fréchet distance between the reference and generated distributions.

### Amino acid classification task

The input to the amino acid classification task is an image containing an amino acid. We created these images in the same way that we created the “first” image from the neighbor location task, as described above, with two differences. First, the images were 11 Å in each dimension, still with 1 Å voxels. Second, the images had a 7th channel, which identified the C_α_of the specific amino acid to classify. To choose this amino acid, we applied several filters to the list of amino acids in the image. First, we randomly dropped amino acids in order give each amino acid type the same probability of being chosen. We determined the empirical distribution of amino acids in the training split of our dataset, which was similar to, but slightly different from, the distribution of all amino acid types in natural sequences (Figure S17). We then dropped each amino acid with probability inversely proportional to this empirical distribution. Second, we dropped the small number of amino acids that had alternate conformations with different amino acid types, as these could not be assigned an unambiguous type. Third, we dropped amino acids for which the sidechain was unlikely to fall completely within the image. To make this determination, we oriented all of the amino acids in our training dataset in a coordinate frame where the C_α_ is at the origin, the backbone N is on the X-axis, the backbone C is in the XY-plane, and the Z-axis is determined using the right-hand rule. In this coordinate frame, we used a dual annealing algorithm [35] to find the sphere, constrained to have a radius of 4 Å, that would contain as many sidechain atoms as possible. The best sphere we found, which we will subsequently refer to as the “sidechain sphere”, was centered at (−1.2729026455200199, −1.9322552322708937, 2.346665604227695) and encompassed 95.5% of the sidechain atoms in our dataset. This coordinate is roughly in the same direction as C_β_, but roughly two carbon-carbon bond lengths from C_α_rather than one. We constructed this sphere for each amino acid in the image, and dropped those amino acids for which the sphere was not entirely contained within the image. After these filtering steps, we uniformly sampled the specific amino acid to classify from those remaining. Finally, we position a sphere with radius 1 Å over the C_α_of the chosen amino acid, then fill in each voxel of the 7^th^ channel with the fraction of that sphere overlapping that voxel. If the image didn’t contain any suitable amino acids, then we left the 7^th^ channel empty.

Our classification models output 21 logits, one corresponding to each amino acid type and one corresponding to the absence of any amino acid. We trained our models using cross entropy loss and the Adam optimizer.

### End-to-end trained model

The first input to the end-to-end training task was a 3D image of a region of macromolecular structure, identical to the input for the diffusion modeling task. The second inputs were crops of that image containing labeled amino acids. Each of these crops was identical to the input for the amino acid classification task, but created in a slightly different way. After performing the filtering steps described above for the amino acid classification task, but on all of the amino acids in the 35 Å diffusion modeling input image rather than just those in an 11 Å image, we uniformly sampled 8 of the remaining amino acids. For each of those amino acids, we uniformly sampled a 11 Å crop that fully encompassed the sidechain sphere used in the initial filtering.

The first part of the training regimen was identical to the diffusion modeling task. We added noise to the 35 Å image and trained a diffusion model to recover the original noise-free image. The second part of the training regimen was identical to the amino acid classification task, except that instead of using a noise-free image as input, we used crops of the image produced by the diffusion model. To combine the mean squared error loss from the diffusion task and the cross entropy loss from the amino acid classification task, we weighted the two losses using learnable parameters, along with a regularization term [30].

The end-to-end model presented in Figure 5 was composed of a diffusion model and an amino acid classification model. The diffusion model was a U-Net [24], composed of a noise embedding, an encoder block, a latent block, and a decoder block. The encoder and latent blocks were SE(3)-equivariant. In these blocks only, we used the convolution, batch normalization, nonlinear, and linear layer implementations provided by the ESCNN library [19] in lieu of the standard implementations. All of the Fourier nonlinearities in this model used the same 96-element grid as described above for the Fréchet distance metric model.

The input to the noise embedding was a single real number giving the standard deviation of the Gaussian noise added to the input image. We converted this number into a 64-dimensional vector using a sinusoidal embedding, with a minimum wavelength of 0.1, a maximum wavelength of 100, and logarithmic spacing. We then used this vector as input to a linear layer with 16-dimensional output, and a ReLU nonlinearity.

The encoder block was composed of a head block and three downsampling blocks. We will refer to the input and output representations of these blocks as *R*(*a*) and *R*(*b*). The head block was identical to that used in the Fréchet distance metric model, described above. The downsampling blocks were similar to the residual blocks used in that same model. They were composed of two branches, followed by a pooling layer. The first branch was composed of a convolution, a batch normalization, a conditioned nonlinearity, a convolution, a batch normalization, and a Fourier nonlinearity. Both convolutions had 3x3x3 kernels, strides of 1, paddings of 1, and no bias. The first convolution had an input representation of *R*(*a*) and an output representation of *R*(*b*). The second convolution had input and output representations of *R*(*b*). The conditioned nonlinearity was composed of a linear layer and a tensor product layer. The linear layer was applied to the noise embedding and transformed it from a trivial input representation to an output representation of *R*(*b*). The tensor product layer was applied to the sum of the input to the conditioned nonlinearity and the transformed noise embedding. The activation function for the Fourier nonlinearity was the first Hermite function. The second branch was composed of a convolution and a batch normalization. The convolution had a 1x1x1 kernel, a stride of 1, a padding of 1, an input representation of *R*(*a*), an output representation of *R*(*b*), and no bias. The sum of these two branches was the input to the pooling layer. The pooling layer was just a con-volution with a 3x3x3 kernel, a stride of 2, a padding of 1, and an output representation of *R*(*b*). The three residual blocks had output representations *R*(4), *R*(8), and *R*(16), with the output from one block being the input to the next. Respectively, these representations had 140, 280, and 560 channels.

The latent block was the same as the downsampling block of the encoder, except for two differences. First, there was no pooling layer. Second, the input and output had the same shape and same representation.

The decoder block was composed of three upsampling blocks and one tail block. We will refer to the number of input and output channels for each of these blocks as *C*_*a*_ and *C*_*b*_, respectively. The up-sampling blocks were composed of an interpolation layer followed by two branches. The interpolation layer increased the size of its input in the spatial dimensions from *n* to 2*n* − 1 via trilinear interpolation. The output of the interpolation layer was summed with the output of the encoder downsampling block of the same shape, forming the residual connections characteristic of the U-Net architecture. This sum was the input to both branches. The first branch was composed of a convolution, a noise block, a batch normalization, a ReLU nonlinearity, a convolution, a batch normalization. Both convolutions had 3x3x3 kernels, strides of 1, and paddings of 1. The first convolution had *C*_*a*_ input channels,*C*_*b*_ output channels, and a bias term. The second convolution had *C*_*b*_ input and output channels, and no bias term. The input to the noise block was the 16-dimensional noise embedding described above. This block was composed of a linear layer, a ReLU nonlinearity, and another linear layer. The linear layers had 64 and 2*C*_*b*_ output channels, respectively. The output from these layers was split into two vectors *M* and *B*, both of dimension *C*_*b*_, then applied to the latent image representation *x* using the formula *Mx* + *B*. Finally, the tail block was composed of a transposed convolution with a 3x3x3 kernel, a stride of 1, and a padding of 0.

The amino acid classification model was composed of an encoder block and a classifier block. The encoder was SE(3)-invariant. In this block, we used the SE(3)-equivariant convolution, batch normalization, and nonlinear layer implementations provided by ESCNN library [19]. In the final step of the encoder, we converted the SE(3)-equivariant latent representation to an SE(3)-invariant vector, with no spatial dimensions. Because the input to the classifier is SE(3)-invariant, the output is too, even though the classifier used standard implementations, not SE(3)-aware implementations, for all of its layers.

The encoder block was composed of a head block, two residual blocks, and an invariant block. The head and residual blocks were both the same as described above for the Fréchet distance metric model, except for the dimensions and representations of the inputs and outputs. The head block had 7 input channels, to account for the extra channel identifying the C^α^of interest, and an output repre-sentation of *R*^′^(7), where 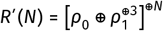, corresponding to 70 channels. The residual blocks had output representations of *R*^′^(14) and *R*^′^(28), respectively, corresponding to 140 and 280 channels. The downsampling layer was only present in the second residual block. The invariant block was composed of a convolution and a Fourier nonlinearity. The convolution had a 3x3x3 kernel and 0 padding. The stride was irrelevant, because the input to this block was also 3x3x3. The input representation for the convolution was *R*^′^(28). The output representation was 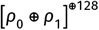, corresponding to 512 channels. The Fourier nonlinearity used a grid of 96 elements from *S*^2^ = *SO*(3)/*SO*(2), i.e. the surface of a unit sphere, used the first Hermite function as the activation function, and had an output representation of 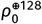, corresponding to 128 channels. The *ρ* representation is SE(3)-invariant, so each element in this 128-dimensional latent vector would be the same regardless of the orientation of the original input image, except insofar as there would be errors relating to the way that different input orientations would be voxelized.

The classifier block was a MLP composed of a first linear layer, a ReLU nonlinearity, a dropout layer, and a second linear layer. The first linear layer had 128 input channels and 64 output channels. The second had 128 input channels and 21 output channels. The dropout layer had a 20% drop probability.

### Sequence recovery

We calculated sequence recovery as follows: (i) create an image of a known protein structure, (ii) create a mask that hides the sidechain atoms in that image but not the backbone or non-protein atoms, (iii) inpaint the masked image using the algorithm described above, (iv) classify each amino acid in the inpainted image, then (v) report the fraction of amino acids that were correctly classified. During training runs for the end-to-end model, we calculated sequence recovery at the end of each epoch using 32 images from the validation dataset. We created the mask in several steps. First, we created a “sidechain image”. We found every sidechain sphere, as defined above but with a radius of 6 Å instead of 4 Å, that could overlap the image. We set each voxel of the sidechain image to the fraction of that voxel that overlapped any of these spheres. Second, we created a “backbone image”. We placed a 2 Å sphere on each backbone C and O atom, and each non-protein atom. We also placed a 0.5 Å sphere on each backbone C^α^and N atom. We chose the former radius to be large enough to prevent the diffusion model from creating new covalent bonds with these atoms, but small enough to allow new H-bonds. We chose the latter radius to be large enough to mask the atom itself, but small enough to allow covalent bonds, since both of these atoms can form covalent bonds with sidechains. As with the sidechain image, we then set each voxel of the backbone image to the fraction of that voxel that overlapped any of these spheres. We did not place spheres on the C^α^atoms, to allow the creation of new C^β^atoms, but note that the C^α^atoms themselves were still included in the backbone image by virtue of being less than 2 Å from the N and C atoms. Third, we combined the sidechain and backbone images using the formula min(*X*, 1 − *Y*), where *X* is the sidechain image and *Y* is the backbone image. When calculating the fraction of amino acids that were correctly classified, we only considered the highest probability prediction made by the classifier, i.e. top-1 accuracy.

## Supporting Information

### SE(3)-equivariant convolutional neural networks

Many of the models discussed in this work are SE(3)-equivariant CNNs. Such models are not widely used in the protein design literature, and the theory behind them is complicated, so the purpose of this section is to provide a gentle introduction. For a more complete treatment, refer to Weiler et al. [31] and Cesa et al. [19].

We will begin by introducing some concepts relating to group theory, the field of mathematics that (among many other things) gives us a language to discuss transformations such as rotations and translations. Each particular kind of transformation is called a group, e.g. the group of 1D translations, the group of 2D rotations with 3-fold symmetry, the group of reflections, etc. Every group is composed of a set of individual transformations, called “group elements”, along with some rule for combining them, called the “group operation”. For example, consider the group of 1D translations. Every 1D translation can be expressed as a real number (and vice versa), so the set of all group elements is ℝ. Applying any two group elements back-to-back gives the same result as applying the sum of those two elements as a single transformation, so the group operation is +. The most general way to identify a group is using the notation (*G*, ⋅), where *G* is the set of group elements and ⋅ is the group operation. Using this notation, the group of 1D translations would be written as (ℝ, +).

There are some common-sense rules that the group elements and operations have to follow. These are known as the “group axioms”. They are:

- There must be an identity element, corresponding to no transformation. For (ℝ, +), the identity element is 0.
- Every element must have an inverse, corresponding to the opposite transformation. Applying an element and its inverse back-to-back must give the same result as applying the identity element, or applying no transformation at all. For (ℝ, +), the inverse of each element *g* ∈ ℝ is −*g*.
- Every combination of group elements must also be a group element. That is, the group operation must be closed with respect to the set of group elements. For a group defined as (*G*, ⋅), this can be expressed as ∀(*a, b*) ∈ *G* ∶ *a* ⋅ *b* ∈ *G*.
- The group operation must be associative. For a group defined as (*G*, ⋅), this can be expressed as ∀(*a, b, c*) ∈ *G* ∶ (*a* ⋅ *b*) ⋅ *c* = *a* ⋅ (*b* ⋅ *c*). For (ℝ, +), this means you would end up in the same place if took a step of 2 + 3 = 5 followed by step of 4, versus a step of 2 followed by a step of 3 + 4 = 7. Note that the group operation is not required to be commutative, so it is not necessarily true that *a* ⋅ *b* = *b* ⋅ *a*.

The group we are most interested in is the group of 3D translations and rotations, as these are the symmetries that are relevant to macromolecules. This group is called *SE*(3): the “special Euclidian” group in 3 dimensions. Here, “special” means that reflections are excluded; the full Euclidean group *E*(3) includes translations, rotations, and reflections, but is not relevant to our work here. *SE*(3) can be broken down into two subgroups: (ℝ^3^, +) for 3D translations, and *SO*(3) for 3D rotations. *SO*(3) is the “special orthogonal” group in 3 dimensions, so named because its elements can be written as orthogonal 3x3 matrices.

All of the groups we’ve mentioned so far—*SE*(3), *SO*(3), (ℝ^3^, +)—have the property that their elements can be expressed as matrices, with matrix multiplication then serving as the group operation. These same matrices can also be used to transform vectors, again using matrix multiplication. The mapping between group elements and matrices is called a “group representation”. To define this term more rigorously, let us begin by noting that the set of all *n* × *n* invertible matrices, together with the matrix multiplication operation, forms a group. This group is called the general linear group of degree *n*, or *GL*(*n*). For any group (*G*, ⋅), a representation is a map *ρ* ∶ *G* → *GL*(*n*) such that *ρ*(*a*⋅*b*) = *ρ*(*a*)*ρ*(*b*) for all *a, b* ∈ *G*. Importantly, representations can be constructed for any *n*. This is what allows us to talk about making 3D rotations to high-dimensional latent vectors buried deep within a neural network, even though we would normally think of such rotations as applying only to 3D vectors.

We can now define “equivariance”. A function is equivariant with respect to some transformation if applying that transformation to the function’s input has the same effect as applying it to the function’s output. This can be expressed as: *f*(*ρ*_in_(*g*)*x*) = *ρ*_out_(*g*)*f*(*x*) for all *g* ∈ *G*, where (*G*, ⋅) is the group of transformations in question, *f* is the equivariant function, *x* ∈ ℝ^*N*^ is the input to that function, *f*(*x*) ∈ ℝ^*M*^ is the output from that function, *ρ*_in_ and *ρ*_out_ are representations of (*G*, ⋅), and *ρ*_in_(*g*) ∈ ℝ^*N*×*N*^ and *ρ*_out_(*g*) ∈ ℝ^*M*×*M*^ are matrices that transform *x* and *f*(*x*), respectively, by *g* ∈ *G*. We’ve previously stated that the advantage of using an *SE*(3)-equivariant neural network is that if it learns to recognize a feature, it will recognize that feature in any location or orientation. We can now say more precisely that, for every layer in that network, if we know what that layer outputs for one location/orientation of a feature and we know what representation that layer is using, then we also know its output for all locations/orientations of that feature.

A similar idea to equivariance is that of “invariance”. A function is invariant with respect to some transformation if applying that transformation to the function’s input has no effect on its output. This can be expressed as: *f*(*ρ*(*g*)*x*) = *f*(*x*) for all *g* ∈ *G*, where these variables are all defined as they were above. Invariance is often a useful property for classification models. For example, a model trained to predict the identity of an amino acid would ideally be *SE*(3)-invariant, so that its predictions would not change if the input was rotated or translated. A simple way to make an *SE*(3)-invariant model is to never give the model any location- or orientation-dependent inputs in the first place. For example, consider a model that requires an atomic structure as input. If we were to encode this structure using only inter-atomic distances, then the model would necessarily be *SE*(3)-invariant, because neither rotating nor translating the structure would change any of the input distances. If we were instead to encode the structure using the 3D coordinates of each atom, then we would need to take additional steps to make the model *SE*(3)-invariant.

We are now prepared to discuss *SE*(3)-equivariant CNNs. In order for any neural network to be equivariant, every layer in the network must itself be equivariant. The three main kinds of layers that comprise a CNN are convolutions, nonlinearities, and batch normalizations. Here we will present just a brief overview of the derivation of an *SE*(3)-equivariant convolution. Refer to Weiler et al. [31] and Cesa et al. [19] for the complete derivation, plus derivations of the other two kinds of layers. The derivation for a convolution begins by establishing two constraints. The first is linearity. That is, we are seeking a function of the form *f*(*x*) = *Wx*, where *x* is a vector and *W* is a matrix that can be learned during training. The second is *SE*(3)-equivariance. We showed above that this constraint can be expressed as *f*(*ρ*_in_(*g*)*x*) = *ρ*_out_(*g*)*f*(*x*), for all *g* ∈ *SE*(3). When we solve for an expression that satisfies both constraints, we find that *f*(*x*) must be a convolution, and its kernel must—loosely speaking—be built from linear combinations of spherical harmonics. This has a few implications. First, it means that *SE*(3)-equivariant convolutions are only negligibly slower than regular 3D convolutions. The only extra step is building the kernel, and this is much faster than the convolution itself (since it’s independent of the size of the input image). Once the kernel is built, there’s no difference between an equivariant and non-equivariant convolution. Second, it means that during training, the optimizer does not directly update the kernel. Instead it updates the coefficients of the linear combinations. And as there are fewer coefficients than there are kernel weights, this significantly reduces the number of learnable parameters. To give a specific numbers, most of the the convolutions in our models used 3x3x3 kernels and representations with some multiple of 35 dimensions. A 3x3x3 convolutional kernel with 35 input channels and 35 output channels (and no bias) would have 3 × 3 × 3 × 35 × 35 = 33075 parameters. An *SE*(3)-equivariant convolutional kernel of the same size, on the other hand, would have only 214 parameters.

Lastly, we will discuss a few practical concerns relating to the design of *SE*(3)-equivariant CNNs. The first is that perfect equivariance is only possible for voxel-aligned transformations. Specifically, 90° rotations and 1-voxel translations. The equivariance error for unaligned transformations isn’t necessarily very significant, though. To minimize this error, it is important to ensure that the training data exhibits all features in all orientations. This is typically accomplished using data augmentation. The second concern is that edge effects arising from padded convolutions can break equivariance. Deep CNNs often use padded convolutions to avoid changing the size of the latent representation after every convolution. However, it’s important to remember that these convolutions work by effectively padding their input with zeros. If a model contains a convolution that is sized such that it uses the padding on one side of its input but not the other, then the input itself will effectively encode its own orientation, and the output of that convolution will no longer be equivariant. The third concern is that pooling layers, if implemented naively, reduce translational equivariance [36]. More specifically, each pooling layer doubles the size of the smallest step that maintains perfect translational equivariance. The solution to this problem is separate the pooling and downsampling operations. The pooling should be done with stride 1, and the downsampling should be done using a Gaussian blur. Furthermore, note that average-pooling maintains rotational equivariance, but max-pooling does not.

## Supplemental Figures

**Figure S1.**
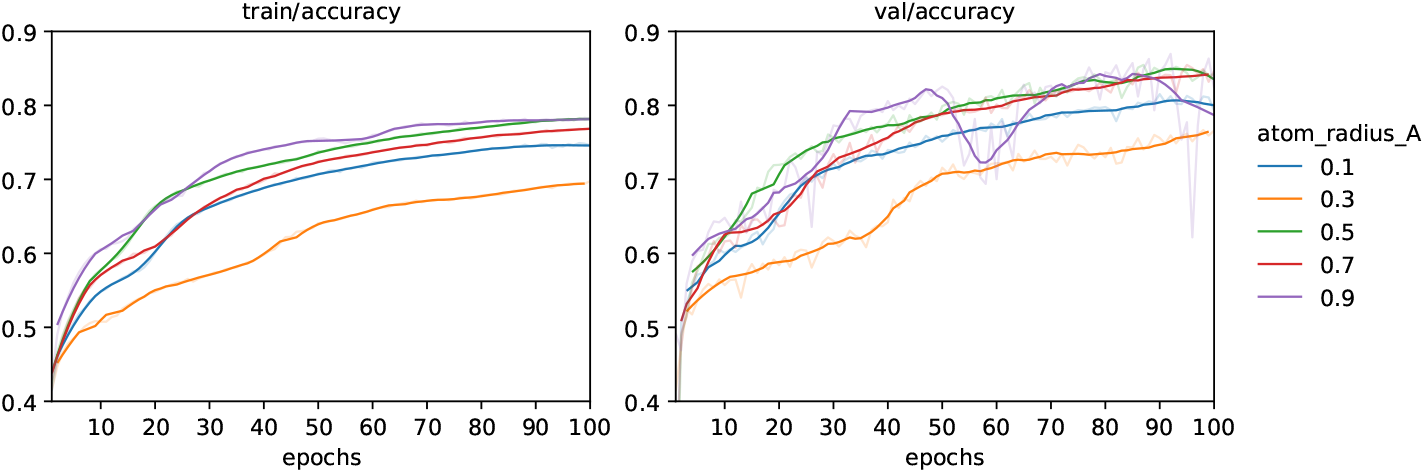
Effect of the atomic radius hyperparameter on the neighbor location pretext task. Faint lines show raw data, bright lines show smoothed curves. The images used in this training run had 1 Å voxels. Note that large atom radii interfere with the ability to create inpainting masks that separate the backbone from the sidechain, and therefore would not be practical even if they performed well.

**Figure S2.**
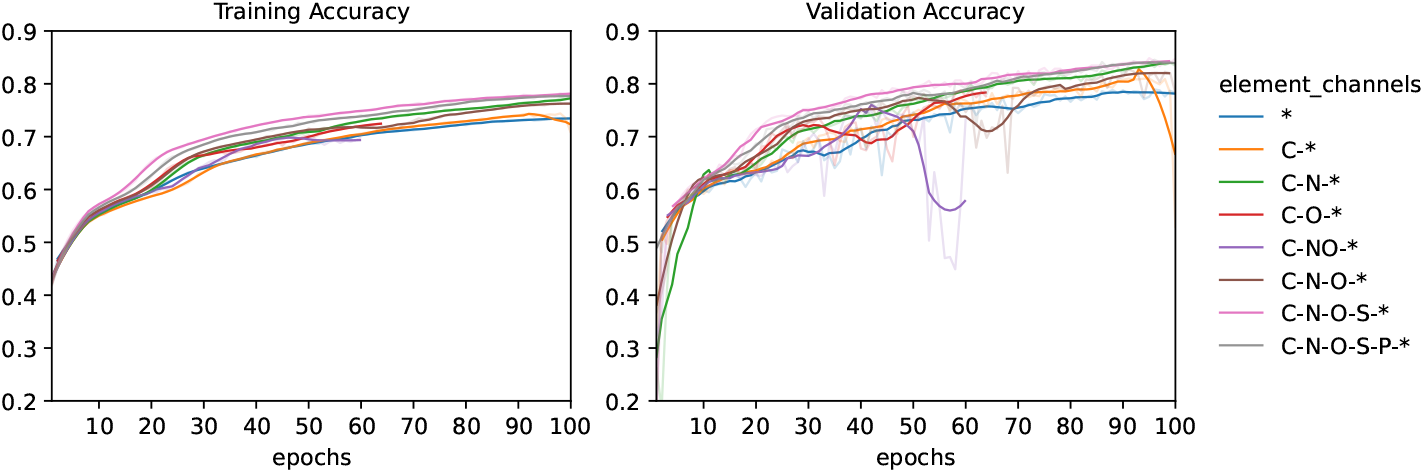
Effect of the element types encoded by the input image on the neighbor location pretext task. Faint lines show raw data, bright lines show smoothed curves. The asterisk means all element types not matched by a previous channel. All of the sulfur channels also include selenium.

**Figure S3.**
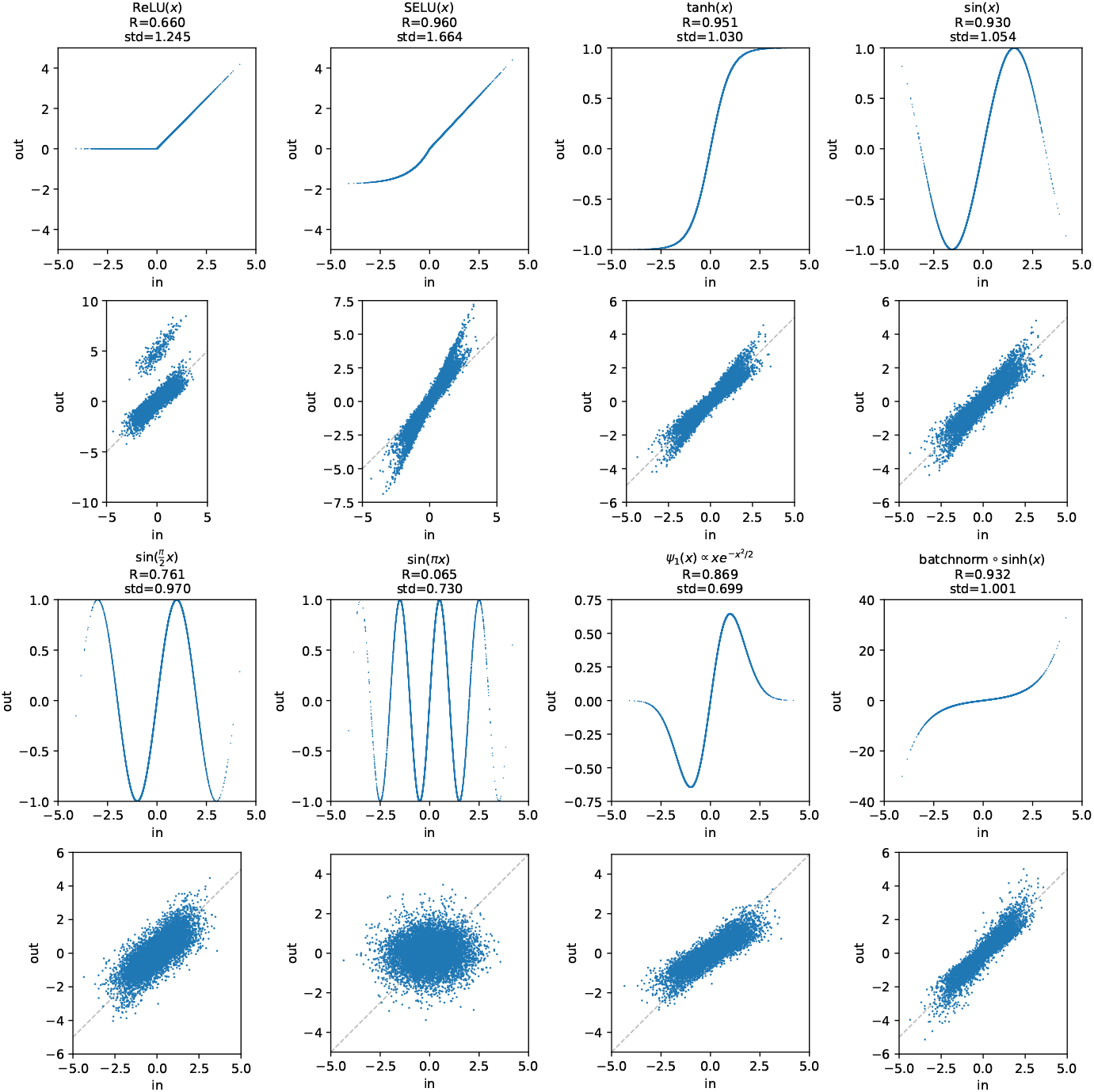
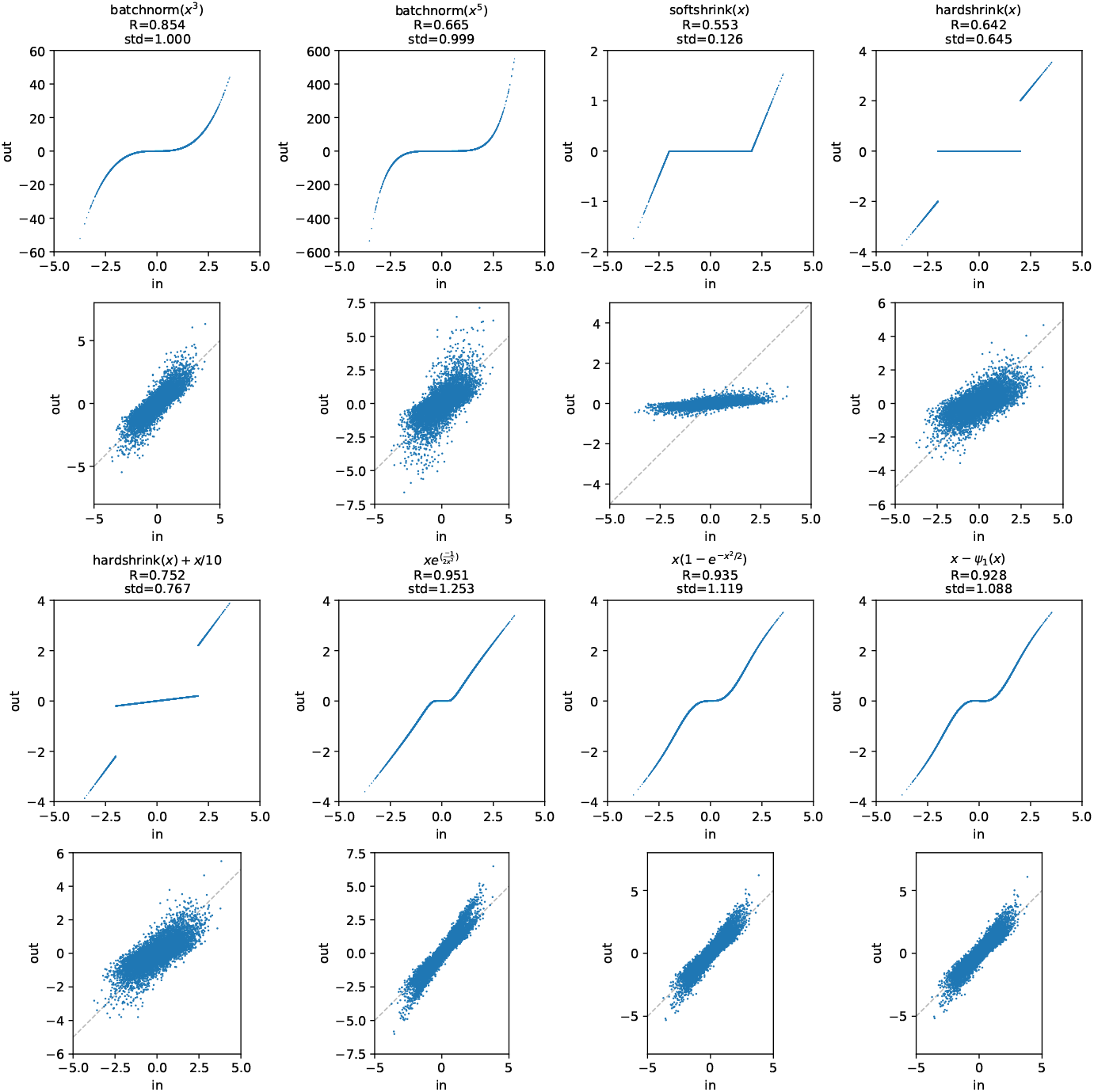
Visualizations of SE(3)-equivariant nonlinear functions. These nonlinear functions are created by performing an inverse Fourier transform, applying the nonlinearity in the spatial domain, then performing a Fourier transform to return to the frequency domain. This is not the only way to create SE(3)-equivariant nonlinear functions, but it one that our models use liberally. Each vertical pair of plots shows a different nonlinearity. In each plot, the x- and y-axes represent the input to and output from the nonlinearity, respectively. The inputs are 9450 points sampled from a Gaussian distribution with a mean of 0 and a standard deviation of 1. This distribution is meant to approximate the inputs to a nonlinear layer following a batch normalization layer. In each pair, the upper and lower plots show the effect of the nonlinearity without and with the Fourier transforms, respectively. The lower plot also includes a grey dashed line with a slope of 1, as a visual reference. The title for each pair lists (i) the correlation coefficient between the input and output values for the lower plot, and (ii) the standard deviation of the output values for the lower plot. Most of the functions we tested were odd, that is: *f*(−*x*) = −*f*(*x*). Functions without this property will exhibit an frequency 0 signal that is shifted above or below the other frequencies. This effect can be seen in the ReLU plot. Most of the models described elsewhere in this work use the first Hermite function, labeled here as *ψ*_1_(*x*), in their nonlinear layers.

**Figure S4.**
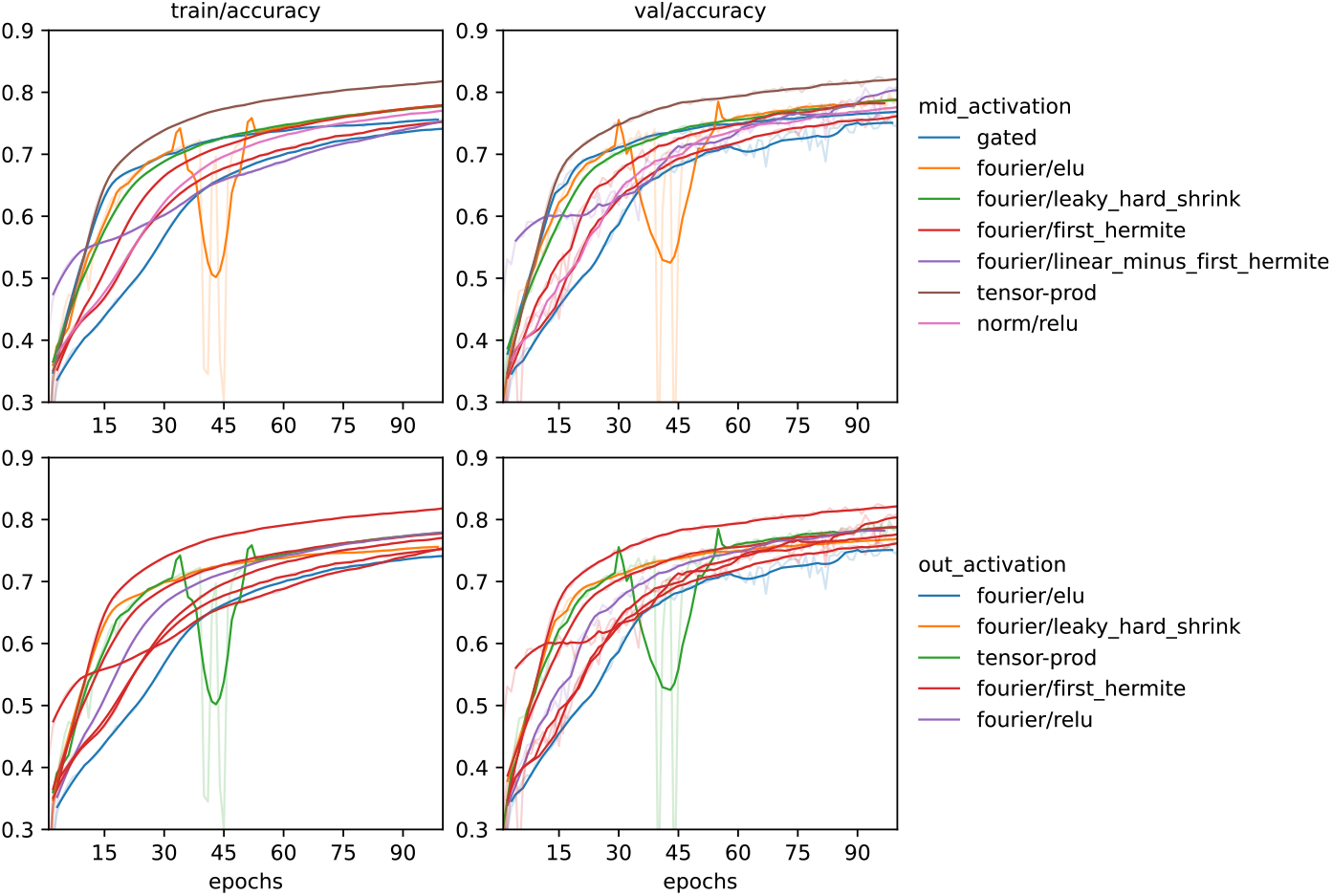
Comparison of select top-performing nonlinear functions on the neighbor location pretext task. Faint lines show raw data, bright lines show smoothed curves. The left column of plots shows accuracy on the training data, while the right shows accuracy of the validation data. Each row of plots shows the same data, but colored according to the a different hyperparameter. The hyperparameters in question are the mid- and out-activations, which refer to the first and second nonlinearities, respectively, present in each residual block. Four different types of nonlinearity are tested here: gated, Fourier, tensor product, and norm. The gated and norm nonlinearities both involve scaling the vectors that make up the latent representation, but differ in how the new magnitudes are calculated. The Fourier nonlinearities interpret the vectors making up the latent representation as signals in the frequency domain, and apply the indicated nonlinear function in the spatial domain. The tensor product nonlinearities are somewhat akin to a polynomial function. For more information, refer to Cesa et al. [19].

**Figure S5.**
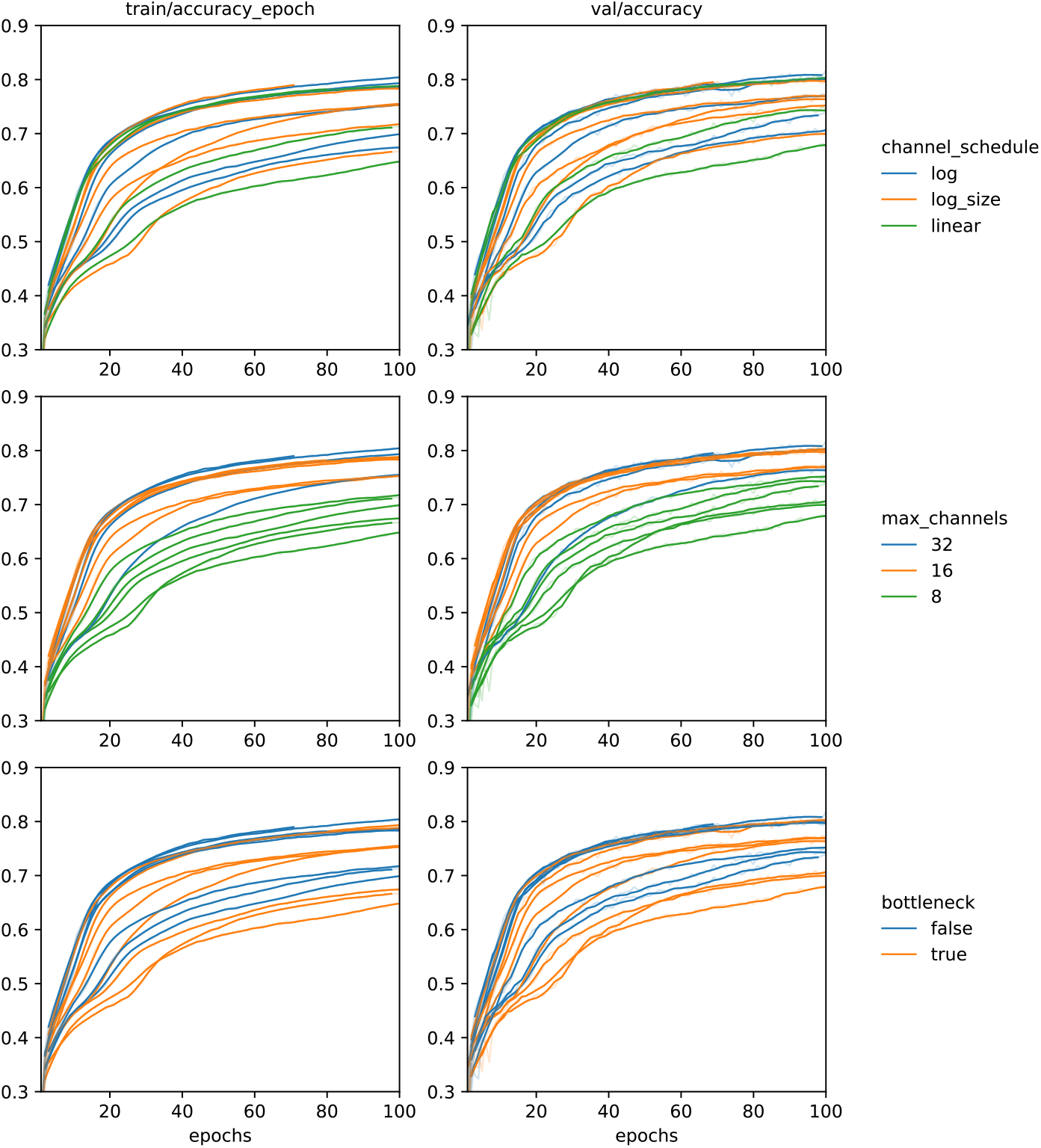
Effect of several hyperparameters related to the width of latent space on the neighbor location pretext task. The left column of plots shows accuracy on the training data, while the right shows accuracy of the validation data. Each row of plots shows the same data, but colored according to a different hyperparameter. The channel schedule determines how the number of channels increases as the depth of the model increases. Refer to the source code for the exact schedules. The “max channels” hyperparameter has a slightly misleading name. It specifies the multiplicity of the representation in the deepest layer of the model. These models all use a 35-dimensional representation,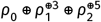 so the actual number of channels is this hyperparameter multiplied by 35. The bottleneck hyperparameter specifies whether or not the number of channels is temporarily halved in the models’ residual blocks.

**Figure S6.**
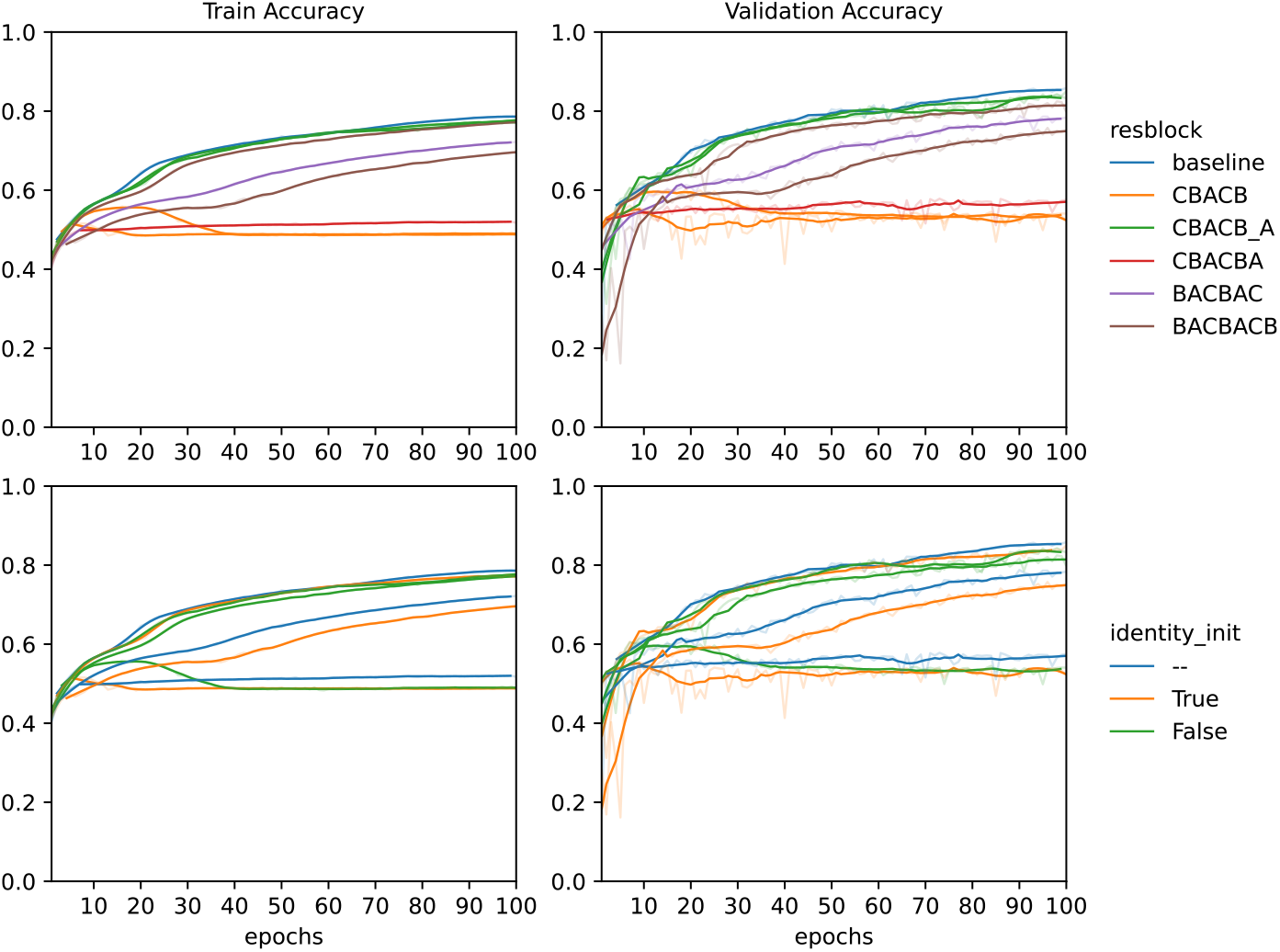
Effect of different residual-block architectures on the neighbor location pretext task. Each architecture is labeled using a string of letters that specify, in order, the layers that are applied before the skip connection. “C” indicates a convolution, “B” indicates a batch normalization, and “A” indicates a nonlinear activation function. The underscore in the “CBACB_A” architecture indicates that the last activation is applied *after* the skip connection. The “identity_init” hyperparameter determines how the weights of the residual block are initialized. If true, they are initialized to make the block behave as an identity function. If false, they are initialized randomly.

**Figure S7.**
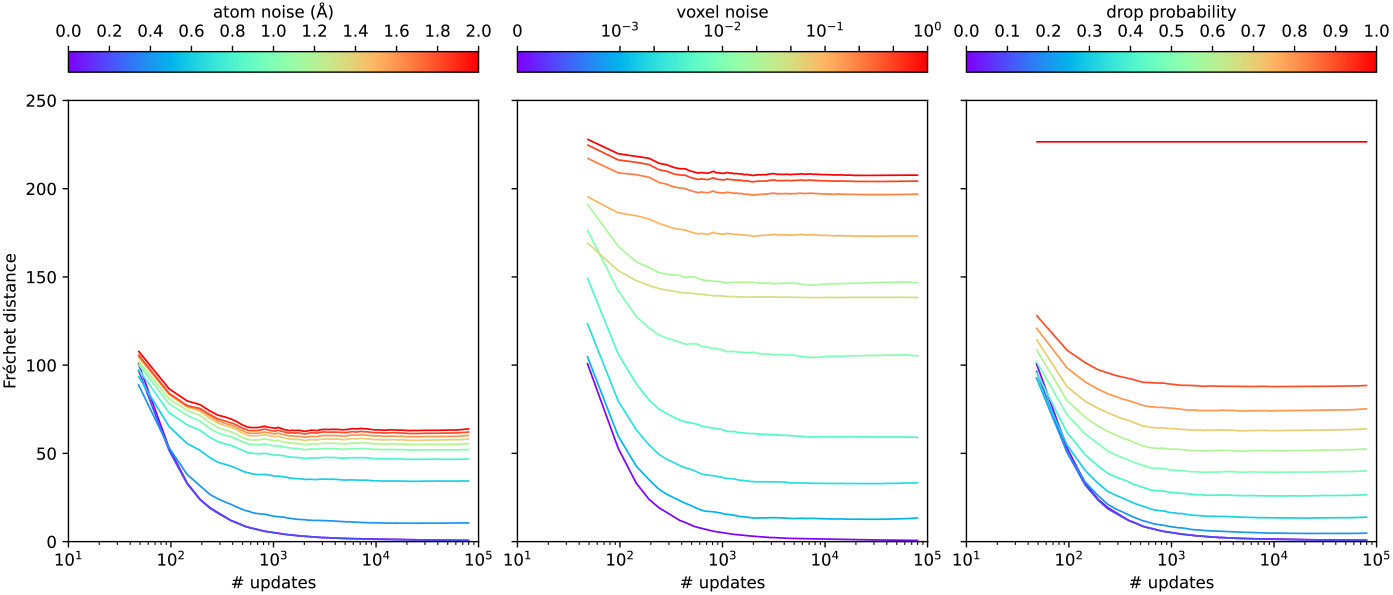
Effects of various perturbations on the Fréchet distance metric. In each plot, the y-axis shows Fréchet distance and the x-axis shows the number of view pairs used to accumulate the statistics underlying the metric. Each 35 Å image contains four non-overlapping view pairs, so the number of images generated is a quarter of the value on this axis. The left plot shows the effect of adding Gaussian noise to the positions of all of the atoms in the structure. The color bar gives the standard deviation of this noise in units of ångströms. The center plot shows the effect of adding Gaussian noise directly to each voxel in the image. The color bar again gives the standard deviation of this noise, but this time the noise is unitless. The right plot shows the effect of randomly removing atoms from the structure. The color bar gives the probability that any particular atom will be removed.

**Figure S8.**
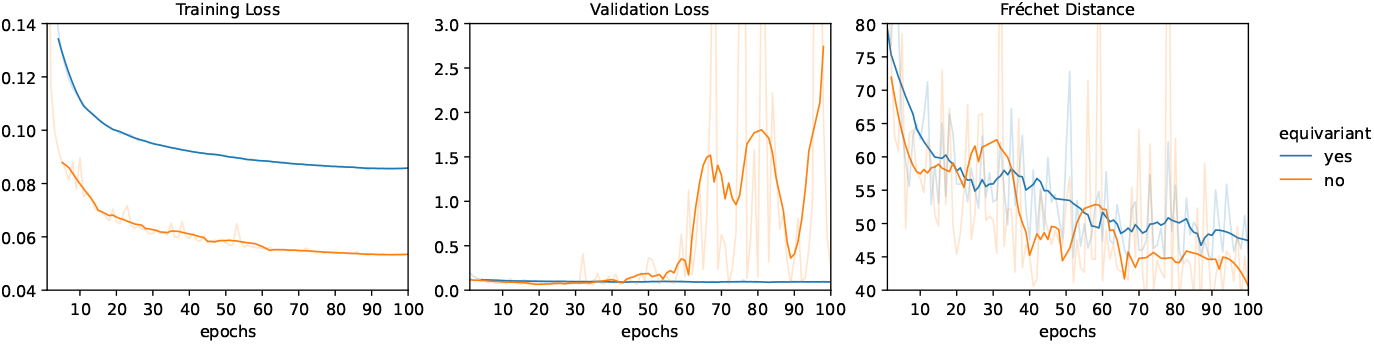
Training a diffusion model with and without SE(3)-equivariance. From left to right, the three plots show training loss, validation loss, and Fréchet distance. Both loss functions are mean-squared error (MSE). Faint lines show raw data, bright lines show smoothed curves. The two models have similarly sized latent spaces. Note that the non-equivariant model achieve better training loss, but severely overfits the training data.

**Figure S9.**
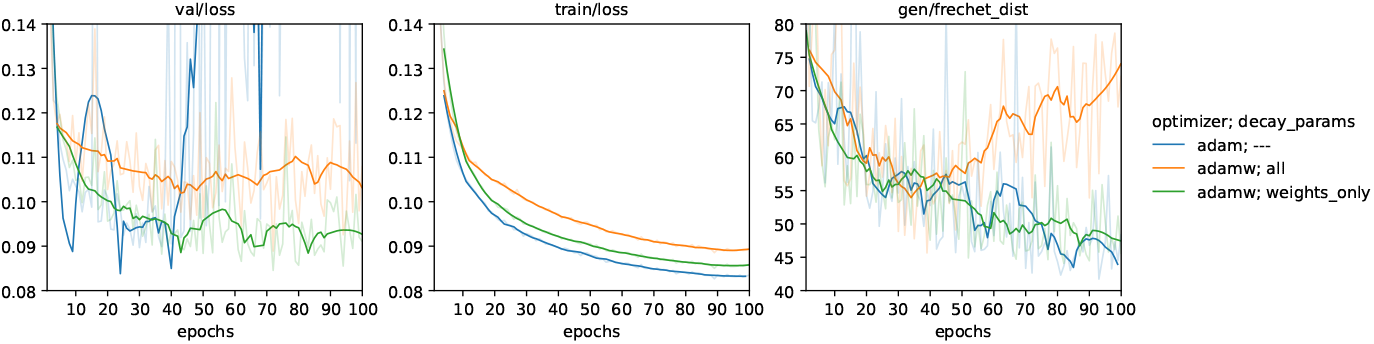
Effect of weight decay diffusion model training. From left to right, the three plots show training loss, validation loss, and Fréchet distance. Both loss functions are mean-squared error (MSE). Faint lines show raw data, bright lines show smoothed curves. The “adam” model was trained without weight decay. The “adamw/all” model was trained with weight decay applied to all parameters. The “adamw/weights_only” model was trained with weight decay only applied to the convolutional kernel weights.

**Figure S10.**
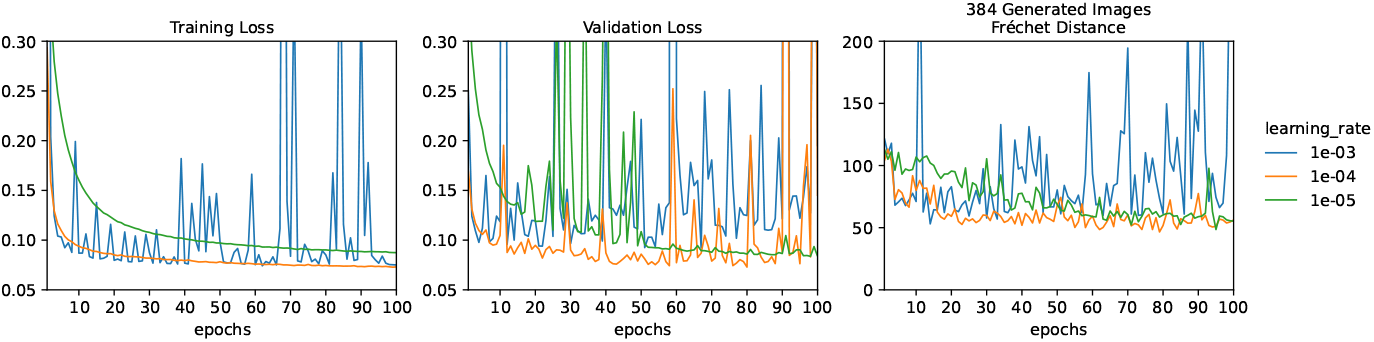
Effect of learning rate on diffusion model training. From left to right, the three plots show training loss, validation loss, and Fréchet distance. Both loss functions are mean-squared error (MSE). With the fastest learning rate (10^−3^), even the training loss does not decrease monotonically. The slower learning rates limit overfitting, but do not eliminate it entirely.

**Figure S11.**
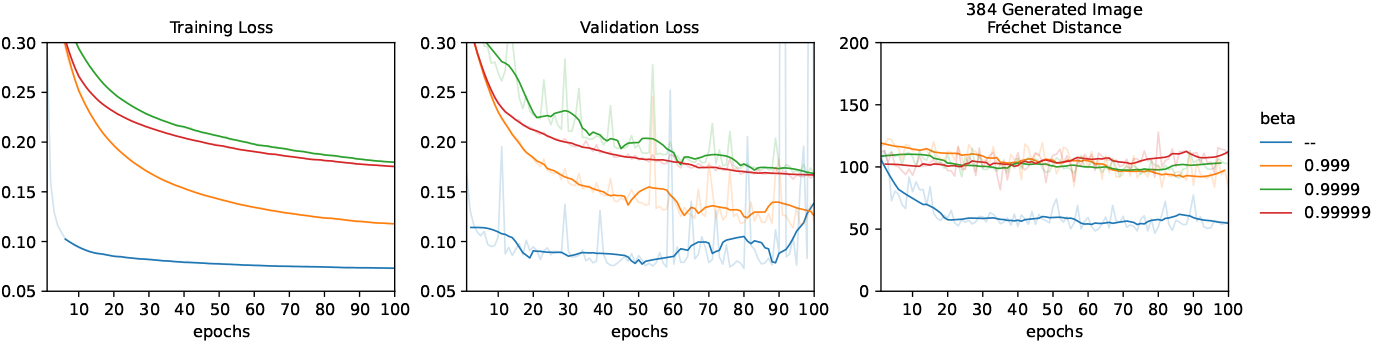
Effect of exponential moving averaging (EMA) on diffusion model training. From left to right, the three plots show training loss, validation loss, and Fréchet distance. Both loss functions are mean-squared error (MSE). Faint lines show raw data, bright lines show smoothed curves. The *β* parameter controls how quickly the moving average forgets old weights, with smaller values forgetting faster. The blue lines have no EMA.

**Figure S12.**
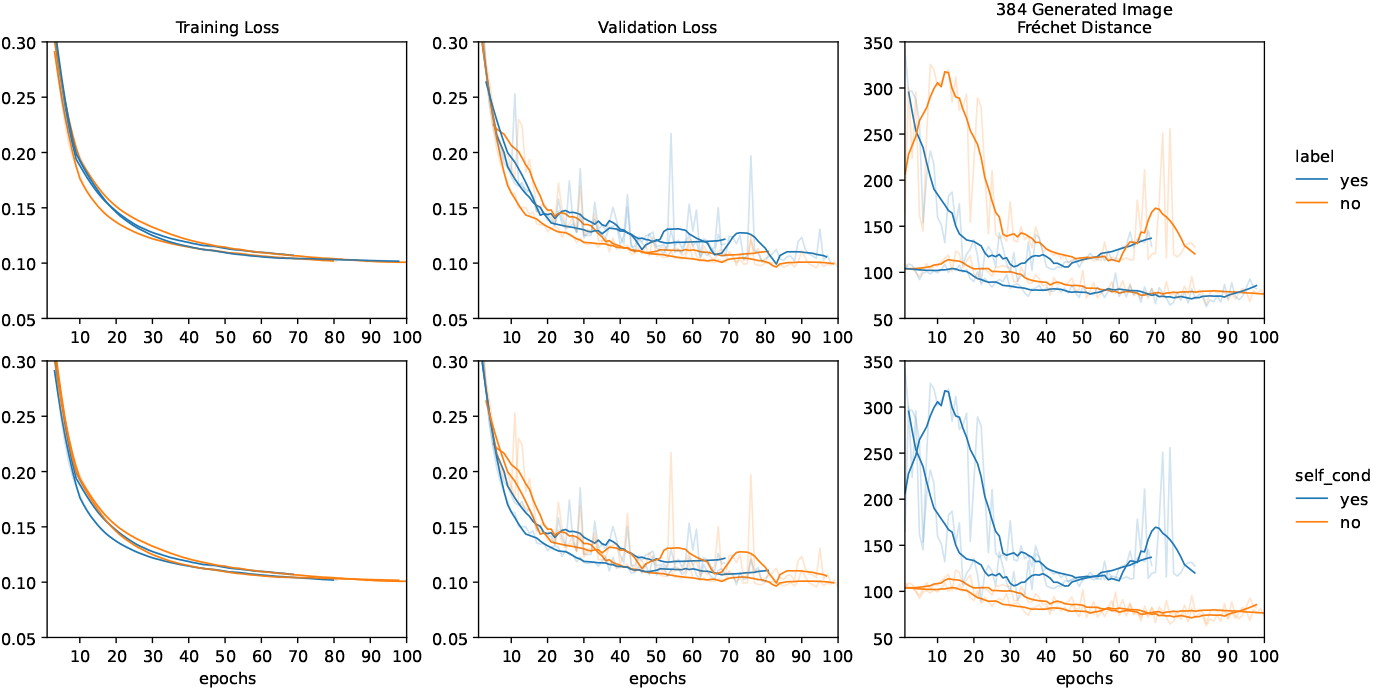
Effect of conditioning strategies on diffusion model training. From left to right, the three columns of plots show training loss, validation loss, and Fréchet distance. Both loss functions are mean-squared error (MSE). Both rows of plots show the same data, but colored according to a different hyperparameter. The hyperparameter in the upper row is whether or not labels encoding molecule type and CATH architecture are provided. The hyperparameter in the lower row is whether or not self-conditioning is employed. Faint lines show raw data, bright lines show smoothed curves.

**Figure S13.**
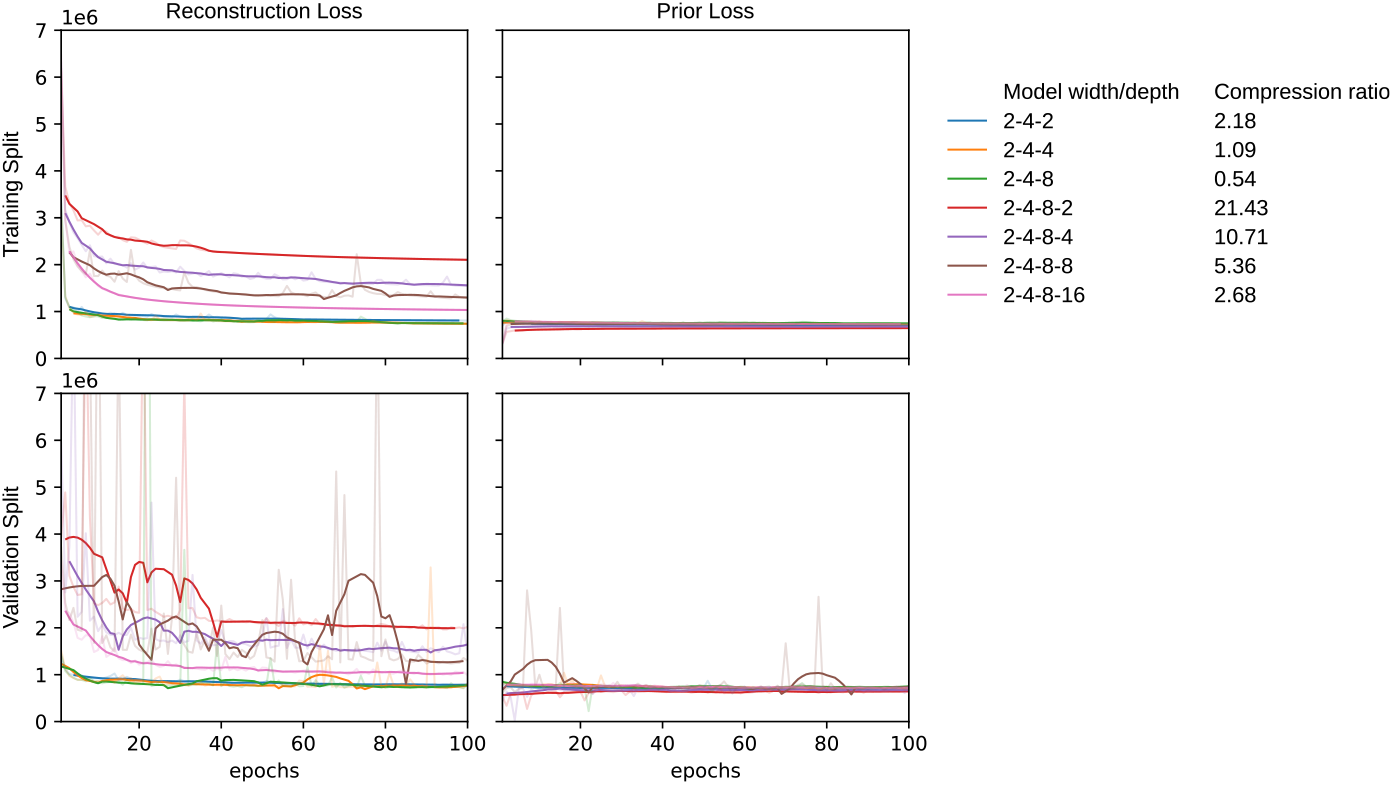
Training a variational autoencoder (VAE) for macromolecular images. The left column of plots shows the reconstruction loss, which is the sum of all differences between the input image and the reconstructed image. The right column shows the prior loss, which is the Kullback–Leibler (KL) divergence between the latent distributions and a standard Gaussian distribution. Note that goal of the actual training run is to minimize the sum of these two loss functions. The top row of plots shows how the models perform on the training split, and likewise the botton row shows the validation split. Turning now to the legend, the model width is a loose description of the dimensions of the latent space at each layer of the model. The compression ratio is the number of voxels in the input image, which is 6 × 35 × 35 × 35 = 257250, divided by the number of voxels in the compressed output image. Note that the models with significant compression ratios (e.g. greater than 5) have poor reconstruction losses.

**Figure S14.**
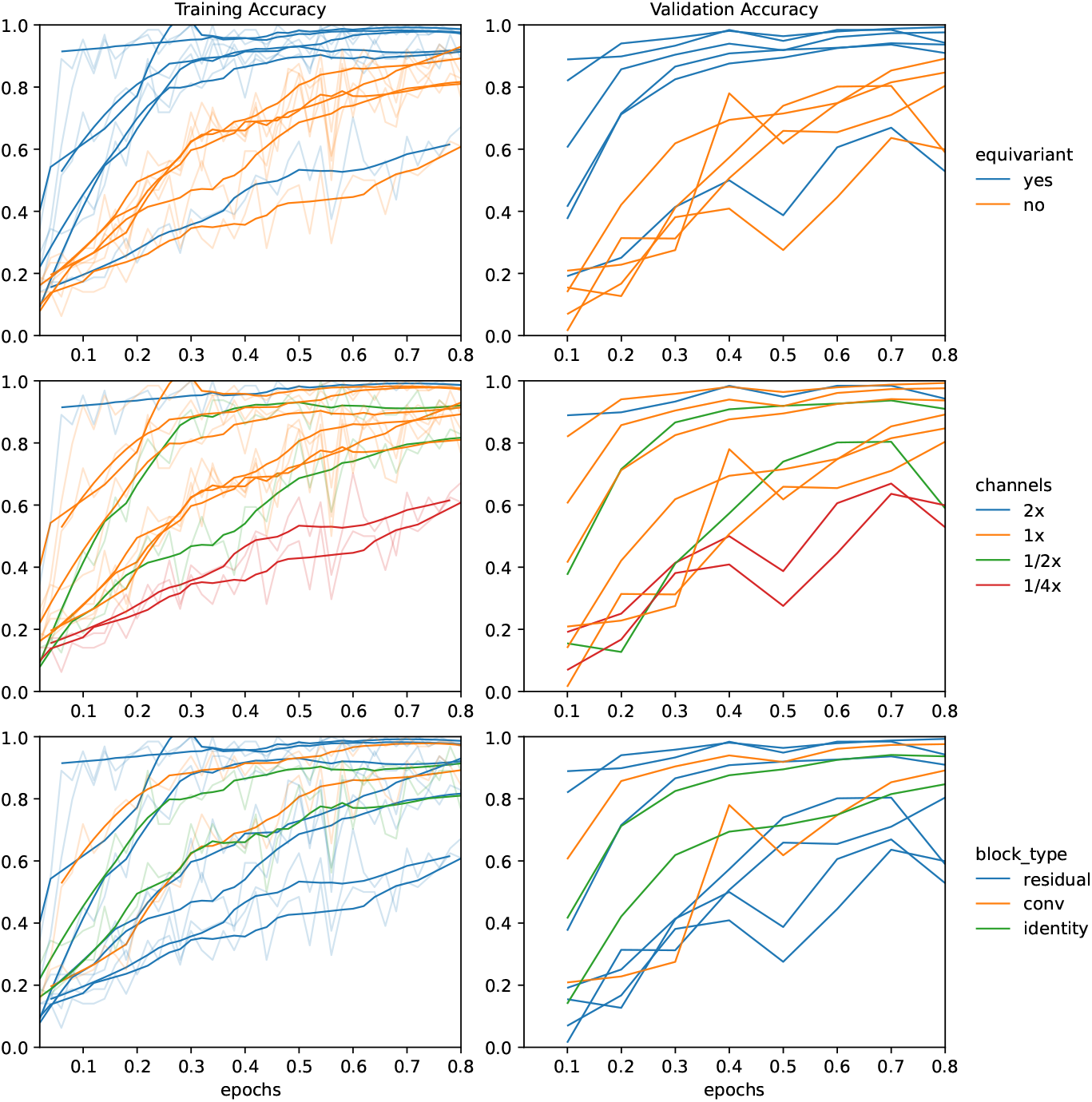
Training amino acid classification models. Faint lines show raw data, bright lines show smoothed curves. The left and right columns of plots show accuracy on the training and validation splits, respectively. Each row of plots shows the same data, but colored according to a different hyperparameter. The “equivariant” hyperparameter indicates whether or not the model is SE(3)-invariant. The “channels” hyperparameter indicates the width of the model, relative to the “1x” baseline. The “block_type” hyperparameter indicates the kind of block that was use to make up the bulk of the model, either “residual” for residual blocks, “conv” for regular convolutions, or “identity” for no blocks other than those used for downsampling. Note that the training runs shown here last for less than 1 epoch; all of these models would perform well after 5-10 epochs. We incorporated the equivariant, 1x, residual model into our end-to-end model. There were simpler models that performed nearly as well, but we (i) preferred a larger model because we expected the end-to-end task to be more difficult and (ii) wanted a model that would be able to update quickly in response to changes in the diffusion model.

**Figure S15.**
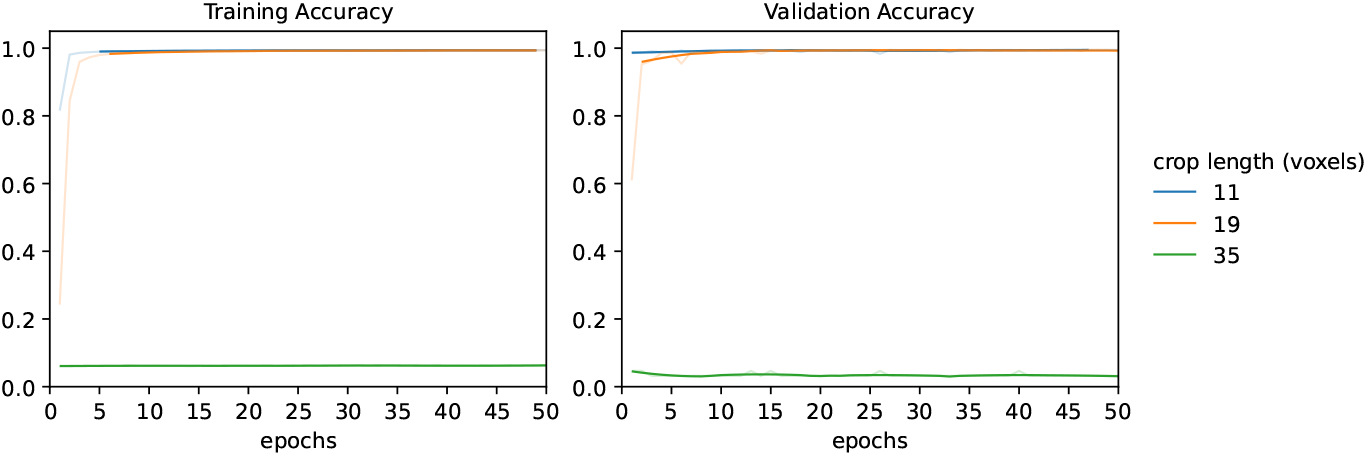
Effect of crop size on the accuracy of amino acid classification models. Faint lines show raw data, bright lines show smoothed curves. The models learn faster with smaller inputs, and are unable to learn anything with inputs that are too big.

**Figure S16.**
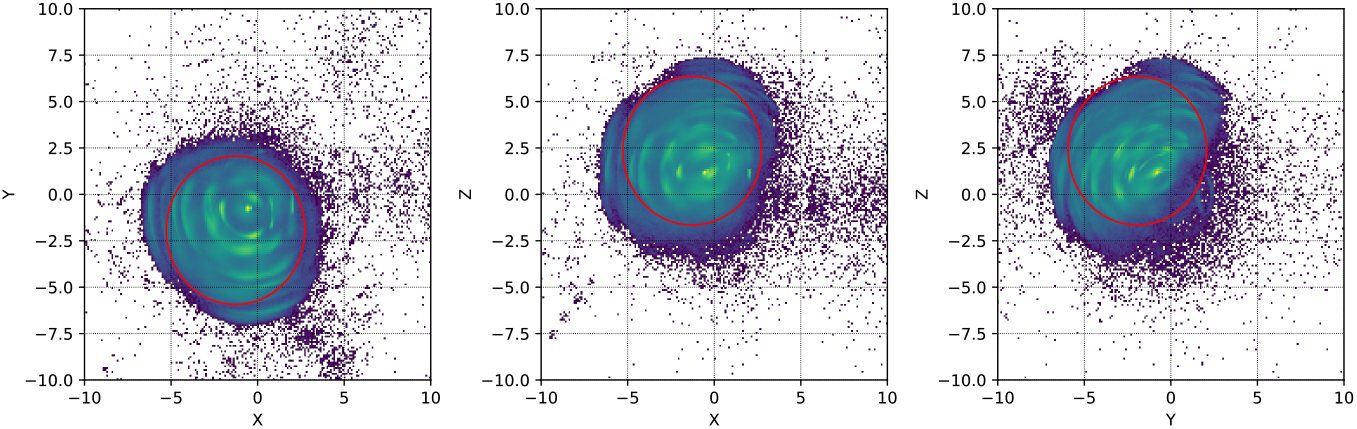
Visualization of the sphere that encompasses >95% of sidechain atoms. Each plot shows the sphere and a sample of sidechain atoms drawn from the PDB projected onto a different 2D plane. The sphere is shown as a red line. The sidechain atoms are shown as a log-scaled heat map where purple/blue bins contain relatively few atoms and green/yellow bins contain relatively many. All of the sidechain atoms are aligned in a coordinate frame defined by the corresponding backbone atoms. All of the axes are in units of ångströms.

**Figure S16.**
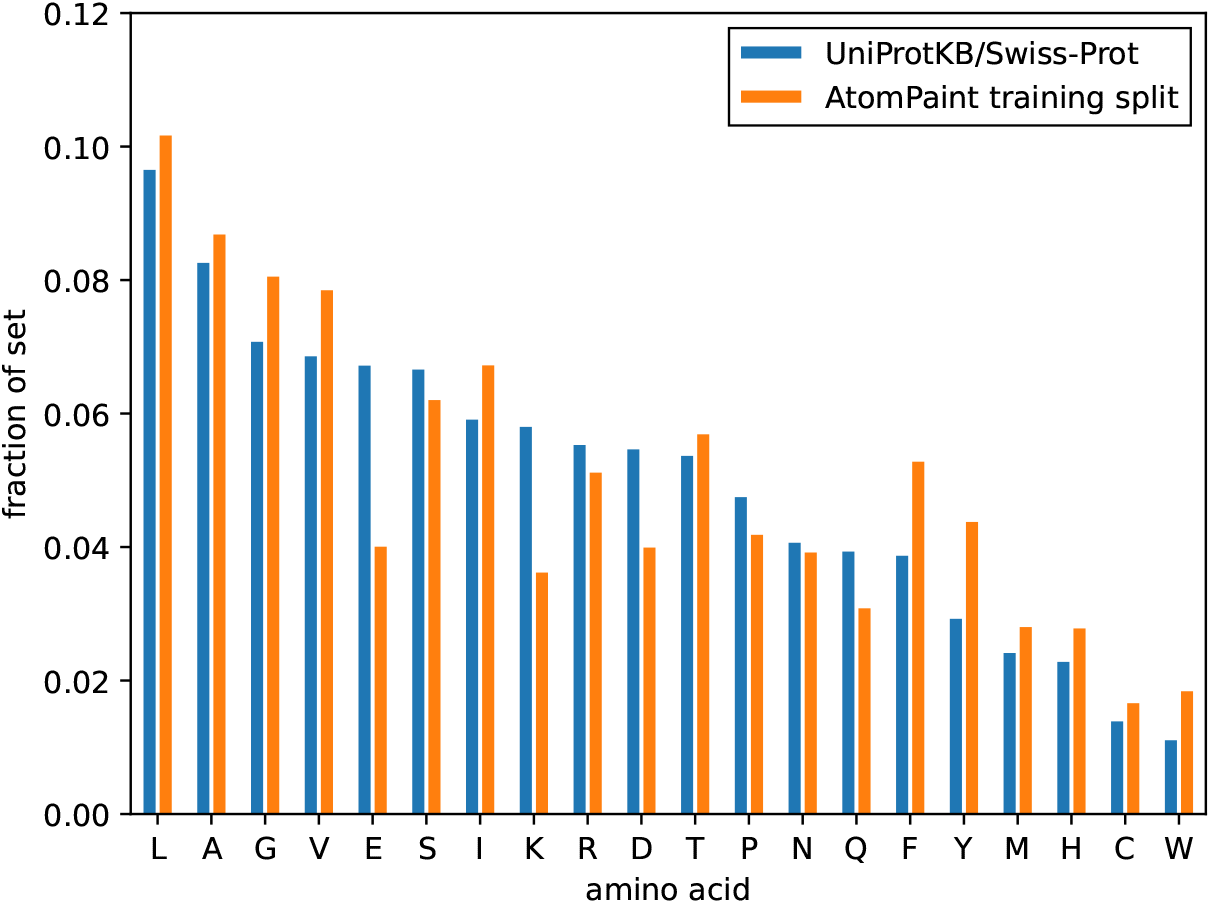
Frequency of each amino acid in (i) the UniProtKB/Swiss-Prot database and (ii) the AtomPaint training
set.

**Table S1.** Sequence recovery predictions made by AtomPaint. 891,086 predictions were made for 527,538 unique residues from 2,617 unique structures sampled from the test split. There are more predictions than there are unique residues because multiple images were inpainted for each structure, and the same residue could appear in more than one of these. The “Examples” column shows how many times each amino acid appeared in the test split. The “Predictions” columns shows how many times each amino acid was predicted by AtomPaint. The “Prediction probability” column converts the prediction counts to probabilities. The “Accuracy” column shows the fraction of the time that AtomPaint makes the correct prediction, conditional on the correct amino acid being the one named in the first column. The “Better than random?” column gives a checkmark to each amino acid for which the accuracy is greater than the prediction frequency, i.e. for which AtomPaint is more accurate than a model that makes random predictions with the same frequencies as AtomPaint.

| Amino acid | Examples | Predictions | Prediction probability | Accuracy | Better than random? |
| --- | --- | --- | --- | --- | --- |
| LEU | 90672 | 23479 | 0.026349 | 0.041457 | ✓ |
| PRO | 36667 | 4623 | 0.005188 | 0.017863 | ✓ |
| VAL | 71024 | 3550 | 0.003984 | 0.006984 | ✓ |
| ALA | 82461 | 25044 | 0.028105 | 0.034998 | ✓ |
| ILE | 60385 | 2419 | 0.002715 | 0.004339 | ✓ |
| GLY | 70021 | 743186 | 0.834023 | 0.920552 | ✓ |
| PHE | 47589 | 1807 | 0.002028 | 0.003005 | ✓ |
| THR | 51315 | 4451 | 0.004995 | 0.006022 | ✓ |
| SER | 53118 | 16251 | 0.018237 | 0.019711 | ✓ |
| ASP | 38087 | 4260 | 0.004781 | 0.006931 | ✓ |
| GLU | 38492 | 6708 | 0.007528 | 0.010496 | ✓ |
| ARG | 39862 | 3087 | 0.003464 | 0.003888 | ✓ |
| ASN | 36474 | 7485 | 0.008400 | 0.009925 | ✓ |
| TYR | 39901 | 395 | 0.000443 | 0.000476 | ✓ |
| HIS | 26016 | 4426 | 0.004967 | 0.006765 | ✓ |
| MET | 24866 | 3075 | 0.003451 | 0.004745 | ✓ |
| GLN | 27355 | 8070 | 0.009056 | 0.011259 | ✓ |
| LYS | 27048 | 27350 | 0.030693 | 0.031832 | ✓ |
| TRP | 16261 | 1196 | 0.001342 | 0.001845 | ✓ |
| CYS | 13472 | 19 | 0.000021 | 0.000000 |  |

## Notes

https://doi.org/10.6084/m9.figshare.30842459

https://doi.org/10.6084/m9.figshare.30826517

### Competing Interest Statement

GMC disclosures: https://arep.med.harvard.edu/gmc/tech.html

### Funding

- Donor Confidential

